# Experimental evidence of stress-induced critical state in schooling fish

**DOI:** 10.1101/2025.02.21.639527

**Authors:** Guozheng Lin, Ramón Escobedo, Xu Li, Tingting Xue, Zhangang Han, Clément Sire, Vishwesha Guttal, Guy Theraulaz

## Abstract

How do animal groups dynamically adjust their collective behavior in response to environmental changes is an open and challenging question. Here, we investigate the mechanisms that allow fish schools to tune their collective state under stress, testing the hypothesis that these systems operate near criticality, a state maximizing sensitivity, responsiveness, and adaptability. We combine experiments and data-driven computational modeling to study how group size and stress influence the collective behavior of rummy-nose tetras (*Hemigrammus rhodostomus*). We quantify the collective state of fish schools using polarization, milling, and cohesion metrics and use a burst-and-coast model to infer the social interaction parameters that drive these behaviors. Our results indicate that group size modulates stress levels, with smaller groups experiencing higher baseline stress, likely due to a reduced social buffering effect. Under stress, fish adjust the strength of their social interactions in a way that leads the group into a critical state, thus enhancing its sensitivity to perturbations and facilitating rapid adaptation. However, large groups require an external stressor to enter the critical regime, whereas small groups are already near this state. Unlike previous studies suggesting that fish adjust their interaction network structure under risk, our results suggest that the intensity of social interactions, rather than network structure, governs collective state transitions. This simpler mechanism reduces cognitive demands while enabling dynamic adaptation. By revealing how stress and group size drive self-organization toward criticality, our study provides fundamental insights into the adaptability of collective biological systems and the emergent properties in animal groups.

## I. INTRODUCTION

Collective behaviors in biological systems have intrigued scientists for decades [1, 2]. These phenomena are observed at all scales of biological organization from macromolecules and cell constituents to groups of organisms and they result from the ability of these biological entities to self-organize and coordinate their actions through their interactions [3–5]. A fundamental question is to understand the adaptive significance of collective behavior in contexts such as environmental changes or to escape dangers [6].

Recently, the concept of criticality derived from statistical mechanics [7] has emerged as a compelling lens through which one can understand the mechanisms underlying adaptability in biological systems. The criticality hypothesis suggests that biological systems should operate at or close to critical points or critical lines that demarcate qualitative changes in collective behaviors and organizations in the phase diagram of parameters, akin to phase transitions in physics [8]. When a biological system is in a critical state, it offers unique properties that maximize its sensitivity to external perturbations, enhance collective information processing and its adaptability in response to environmental cues [9–11].

Recent empirical evidence has begun to shed light on the relevance of criticality in various biological systems. From neural dynamics [12–15] to gene regulation, aggregation in social amoeba [16], and collective motion in animal groups [17, 18], studies have reported instances of scale invariance and critical behavior. In particular, research on spontaneous behavioral cascades and escape waves in schooling fish has provided valuable insights into how these systems could operate near criticality [19, 20]. Fish schools could also modulate the distance to criticality according to the environmental context. In groups of juvenile golden shiners (*Notemigonus crysoleucas*), it has been suggested that when the perceived risk increases, the distance from criticality decreases [21]. Fish schools would thus be able to achieve a fine balance between their sensitivity and the robustness of their collective responses as a function of the riskiness and noisiness of the environmental conditions. In that context, a crucial open question is that of the actual mechanisms used by these animal collectives to tune themselves toward the critical region of the parameter space [22].

Here, we address this question through the study of collective phase changes in schooling fish in response to a stressing condition. We combine experiments with computational modeling to investigate how group size and stress affect the way fish swim and interact with each other, and the consequences on the collective behavior exhibited by groups of rummy-nose tetras (*Hemigrammus rhodostomus*). Our results suggest that the stress state of individuals is strongly modulated by group size, presumably due to a social buffering effect [23, 24]. Moreover, when the fish are stressed, they spontaneously adjust the intensity of their social interactions such that the shoal collectively reaches the proximity of a critical state, which allows it to rapidly adapt its behavior in response to changes occurring in the environment.

## II. RESULTS

### A. Effect of mild stress on collective swimming behavior

The experiment consists of inducing mild stress in a group of fish by abruptly increasing the intensity of light from 0.5 lx to 25 lx. Each of our experiment begins with a 10-minute habituation period at 0.5 lx. A sudden change in the light intensity is then repeated 10 times, involving a series of 2-minute intervals at 0.5 lx alternating with 1 minute intervals at 25 lx (see Fig. 1). This latter condition, called the *Stressing condition*, is replicated 6 times with groups of *N* = 10 and *N* = 25 fish (see Supplemental Material (SM) Movie S1 and S2 [25]). In the *Control condition*, the light intensity is kept to a constant value of 25 lx. Here, too, each experiment begins with a 10-minute habituation period and the fish behavior is then recorded for one hour with no changes in light intensity. The Control condition is replicated 4 times with groups of *N* = 10 and *N* = 25 fish (see SM Movie S3 and S4 [25]). Details on experimental protocol are provided in section IV.A.

**FIG. 1.**
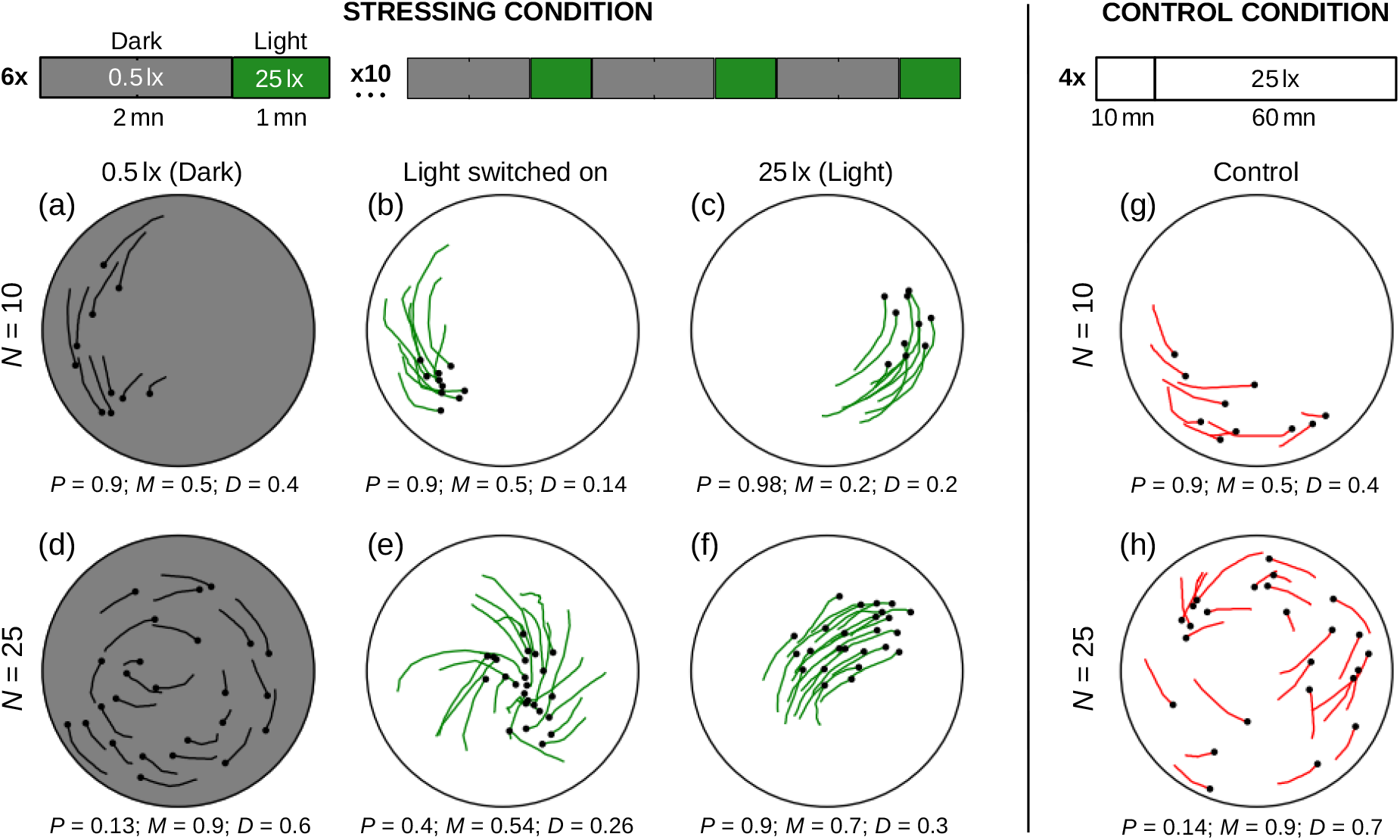
Representative states of fish groups in Stressing and Control conditions. (a–f) Stressing condition: 6 replicates of 10 periods of 3 minutes for each group size alternating low light intensity (Dark, 0.5 lx during 2 min) and high light intensity (Light, 25 lx during 1 min). (a, d) Snapshots of a group of *N* = 10 and *N* = 25 right before the increase in light intensity, (b, e) right after, and (c, f) 30 s later. (g, h) Control condition: 4 replicates of 70 min at constant high light intensity (25 lx) for each group size. Snapshots of a group of *N* = 10 and *N* = 25 in the Control condition. Black dots represent fish positions and the black, green, and red lines are the individual trajectories over the past 1 s. The instantaneous values of polarization *P*(*t*), milling *M*(*t*), and dispersion *D*(*t*) are provided for each spatial snapshot.

Previous studies have shown that when fish are subjected to a sudden increase in light, their physiological response is very similar to that induced by a threat and resulted in a very high increase in oxygen consumption [26]. Here, as depicted in Fig. 1(a-f), sudden changes in light conditions induce qualitative shifts in the collective motion of fish groups. We quantify collective motion via three group-level instantaneous observables: polarization *P*(*t*), which measures the degree of alignment of fish; milling *M*(*t*), which measures how much the fish turn in the same direction around the center of the tank, independently of the direction of rotation; and dispersion *D* (*t*), which measures the spread of the group. The three observables take dimensionless values in [0, 1] and their mathematical expressions are given in section IV.B. When the intensity of light increases, the fish immediately move towards each other while remaining closer to each other, leading to a decrease in the dispersion ⟨*D*⟩ (*t*) from 0.42 at 0.5 lx to 0.22 in groups of 10 fish [Fig. 2(c)], and from 0.64 to 0.4 in groups of 25 fish [Fig. 2(f)]. Then, during the next 60 s at 25 lx, the fish disperse slightly again, but still remain relatively more cohesive than at 0.5 lx or in the Control condition [red solid lines, Fig. 2(c, f)]. In both group sizes, the high cohesion, combined with the propensity to swim near the wall, gives rise to higher polarization [Fig. 2(a, d)] and smaller milling [Fig. 2(b, e)]. This effect is more pronounced in large groups: ⟨*P*⟩ (*t*) grows from 0.65 to 0.83 in small groups [Fig. 2(a)] and from 0.25 to 0.55 in large groups [Fig. 2(d)], while ⟨*M*⟩ (*t*) decreases from 0.53 to 0.40 in small groups [Fig. 2(b)] and from 0.73 to 0.49 in large groups [Fig. 2(e)].

**FIG. 2.**
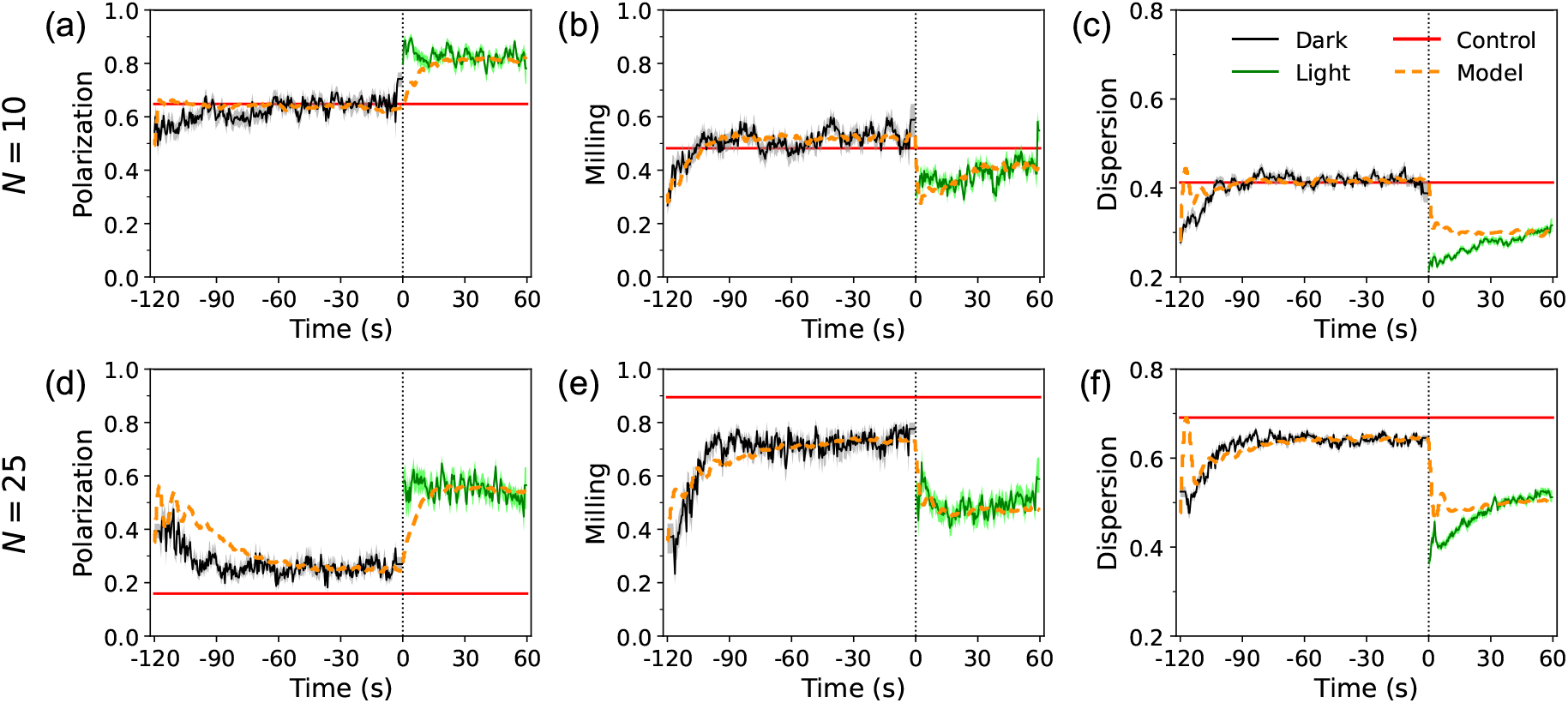
Temporal dynamics of collective observables. Time series of polarization, milling, and dispersion, averaged over repeated conditions, in the Dark (black lines) and Light (green lines) periods of the Stressing condition, for *N* = 10 and *N* = 25 fish. Solid lines represent the experimental results, red lines the mean value of the Control condition, and orange dashed lines the numerical simulations of the model. The shaded areas represent the 68% confidence interval (bootstrap of 1000 rounds) for the experimental results. The vertical black dotted line indicates the time at which the light is switched on. Mean *±* standard deviation from the last 30 seconds of Dark condition to the last 30 seconds of Light condition for *N* = 10: *P* = 0.65 *±* 0.22 to 0.83 *±* 0.18, *M* = 0.53 *±* 0.26 to 0.40 *±* 0.22, *D* = 0.42 *±* 0.08 to 0.28 *±* 0.06. And for *N* = 25: *P* = 0.25 *±* 0.16 to 0.55 *±* 0.24, *M* = 0.73 *±* 0.19 to 0.49 *±* 0.25, *D* = 0.64 *±* 0.04 to 0.49 *±* 0.07.

Then, when the light intensity drops again, the fish disperse and recover in less than 30 s the state they initially had in low light intensity. Fig. 2(c, f) show that dispersion quickly recovers a constant “rest” value (exactly the control value when *N* = 10, and close to the control value when *N* = 25) with a relaxation time of about 10 s in both group sizes. This contrasts with the dynamics observed after the increase in light intensity, in which the relaxation time of the dispersion is much longer in both group sizes (over 60 s). Altogether, these results suggest that the increase in light intensity induces a change in the behavioral state of fish which is not simply due to the higher cohesion of fish at 25 lx, but a consequence of the stress induced by the sudden change in light condition.

To accurately compare the difference between the state of fish at 25 lx having experienced an abrupt change in light condition and the state of fish that have been exposed to a constant light intensity of 25 lx, we computed the probability density functions (PDF) of polarization, milling, and dispersion in the Stressing (black and green solid lines in Fig. 3) and Control (red solid lines in Fig. 3) conditions. With the exception of milling in small groups [Fig. 3(b)], the three measures exhibit clear differences between the Light period of the Stressing condition and the Control condition for both group sizes, while the values observed in the Dark period of the Stressing condition and in the Control condition are relatively similar.

**FIG. 3.**
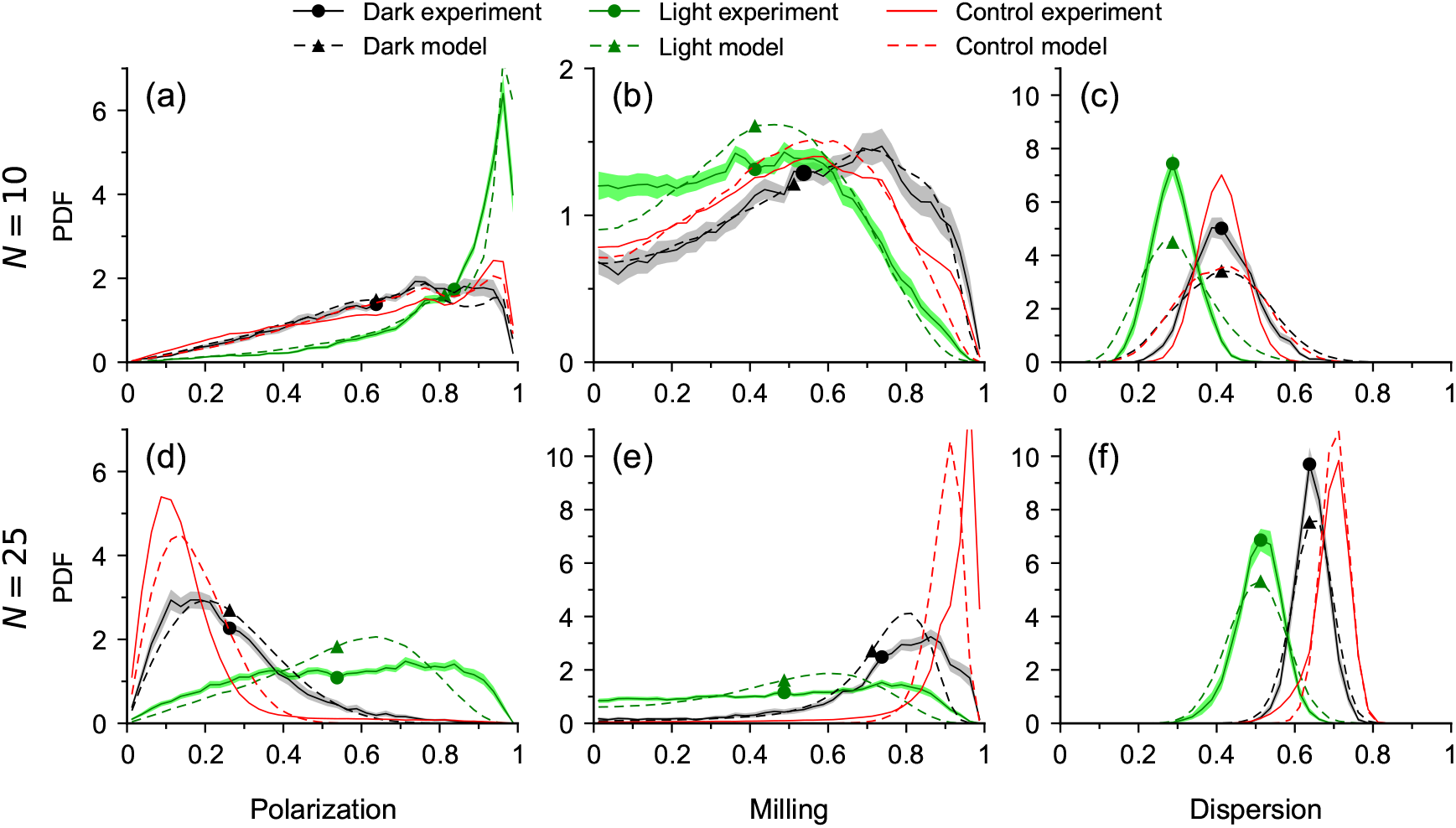
Distributions of collective observables. Probability density function (PDF) of polarization, milling, and dispersion observed in groups of *N* = 10 fish (a–c) and *N* = 25 fish (d–f) in the Stressing (dark and green solid lines) and Control (red solid lines) conditions, together with the results of the numerical simulations of the model (dashed lines). The shaded areas represent the 68% confidence interval (bootstrap of 1000 rounds) for the experimental results. Colored symbols represent mean values also reported in SM Table S2 [25].

Dispersion is smaller in the Light period of the Stressing condition than in the Control condition [Fig. 3(c, f)] for both groups. Polarization and milling, however, show some group size dependence. In small groups, the polarization is high with a mode at *P ≈* 0.95 [Fig. 3(a)], meaning that fish are very often extremely well aligned, much more than what was suggested by the mean value in Fig. 3(a). The small bump visible at *P ≈* 0.8 in Fig. 3(a) corresponds to the situation where all the fish of the group except one are directionally aligned (*P ≈* (9 ™ 1)*/*10 = 0.8). In the Dark period of the Stressing condition [black solid line in Fig. 3(a)] and in the Control condition [red solid line in Fig. 3(a)], the PDF is more uniformly distributed, and where the small bump at *P ≈* 0.8 is still visible. In small groups, the PDF of the milling in the Light period of the Stressing condition is similar to the one observed in the Control or Dark period of the Stressing condition [Fig. 3(b)], contrary to what is observed with the polarization [Fig. 3(a)].

In large groups, the differences between the Light period of the Stressing condition and the Control condition are much more evident. In the Control condition, polarization is weak ⟨*P*⟩ = 0.16 with a mode at *P ≈* 0.1 [see Table S2 [25] and red solid line in Fig. 3(d)], a value which is comparable to the mean polarization of 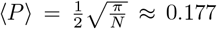 expected assuming uncorrelated random orientations of fish. Milling is also quite high, with ⟨*M*⟩ = 0.90 and peaked at *M ≈* 0.95 [see Table S2 [25] and red solid line in Fig. 3(e)]. Low polarization ⟨*P*⟩ = 0.25 and high milling ⟨*M*⟩ = 0.74 [see Table S2 [25] and black solid lines in Fig. 3(d, e)] are also observed in the Dark period of the Stressing condition. These PDFs together with the visual inspection of the videos show that both in the Control and in the Dark period of the Stressing conditions, the fish rotate along the edge of the tank, with the same direction of rotation leading to a high value of *M*, and consequently, a small value of *P*. By contrast, in the Light period of the Stressing condition, the PDFs of polarization and milling are both almost uniform [green solid lines in Fig. 3(d, e)].

In the next section, we use a data-driven behavioral model to unveil the mechanisms by which groups of fish react to the Stressing condition and how these observed dynamics reflect the proximity to a critical region.

### B. Modelling collective response of fish school to intermittent light condition

We use the burst-and-coast model introduced in [27] that describes the swimming mode and social interactions of *H. rhodostomus* also interacting with the wall of the circular tank. This model has been extended in [28] to account for the modulation of social interactions between fish and their reactions to obstacles by light intensity (ranging from 0.5 lx to 50 lx) and its impact on collective motion patterns in groups of different sizes, spanning the range *N* = 1 to 25. Our broad strategy is as follows: We first parametrize the values of movement and social interactions that shape the collective movement of fish schools in Control, Dark and Light conditions. We then construct the phase diagram of the model and ask if the estimated parameters lie in the vicinity of transitions between different collective motion phases.

The model is based on the assumption that the behavior of a fish results from the combination of three factors: the spontaneous behavior of fish characterized by stochastic spontaneous heading changes of amplitude *γ*_R_; the repulsion of the wall of the tank, of intensity *γ*_w_; and the effect of social interactions, consisting of long-range attraction/short-range repulsion and alignment interactions, whose strength are denoted *γ*_Att_ and *γ*_Ali_, respectively. Moreover, based on previous empirical evidence, the model considers that at each time, each fish only interacts with its two most influential neighbors and combines their influence in an additive form (see Section IV.C for a detailed description of the model). The model simulates the behavior of *N* fish under each light condition and provides the individual trajectory of each fish.

The numerical simulations carried out in [28] show that fish adapt their swimming behavior and social interactions to each light condition ranging from 0.5 lx to 25 lx, and for different group sizes spanning from *N* = 1 to 25, essentially through the modulation of the three parameters *γ*_R_, *γ*_Att_, and *γ*_Ali_, while the other parameters remain unchanged. We therefore explore the 3D parameter space of (*γ*_R_, *γ*_Att_, *γ*_Ali_) to find the values for which the model best reproduces the experimental results obtained in the three lighting conditions. We have kept the parameter values that remain unchanged equal to those determined in [28]. The result is a triplet of values ***γ*** = (*γ*_R_, *γ*_Att_, *γ*_Ali_) for each light condition and each group size (see Table I). The procedure used to determine the parameter values in the three lighting conditions is described in detail in Section IV.D.

**TABLE 1.** Estimation of parameter values of the noise (*γ*_R_), the strength of attraction (*γ*_Att_), and the strength of alignment (*γ*_Ali_) in the Control and Stressing conditions (Dark and Light periods) and for each group size. In the Dark and Light periods, the values have been estimated 30 s after the change of light intensity. In the Stressing condition, arrows indicate the variation of the parameter values after the change of light intensity.

| | $N = 10$ | | | $N = 25$ | | |
| --- | --- | --- | --- | --- | --- | --- |
|  | Control | Dark | Light | Control | Dark | Light |
| $\gamma_R$ | 0.24 | 0.2 | ↘ 0.04 | 0.04 | 0.16 | ↗ 0.34 |
| $\gamma_{Att}$ | 0.023 | 0.026 | ↗ 0.034 | 0.013 | 0.018 | → 0.016 |
| $\gamma_{Ali}$ | 0.022 | 0.023 | → 0.025 | 0.004 | 0.007 | ↗ 0.021 |

In the Stressing condition, the sudden increase in light intensity leads to a change in the strength of interactions between fish. However, this effect also depends on the size of the group. In groups of *N* = 10, fish react by increasing the attraction strength, while in groups of *N* = 25, fish increase the alignment strength. At the same time, the spontaneous heading changes of fish *γ*_R_ decrease in groups of *N* = 10 fish, but increase in groups of *N* = 25 fish. These results suggest that the impact of stress on fish depends on the size of the group. With the exception of *γ*_R_ in groups of *N* = 25 fish, the parameter values between the Control and the Dark conditions are comparable. In both group sizes, the intensity of social interactions is similar, but the amplitude of the spontaneous heading changes of fish is much smaller in the Control condition than in the Dark in groups of *N* = 25 fish.

To simulate the impact of Stressing condition on individual behavior, we consider that the values of parameters *γ*_Att_ and *γ*_Ali_ increase abruptly when the light is turned on, and then slowly decrease to reach a constant value after 30 s (Fig. 4).

**FIG. 4.**
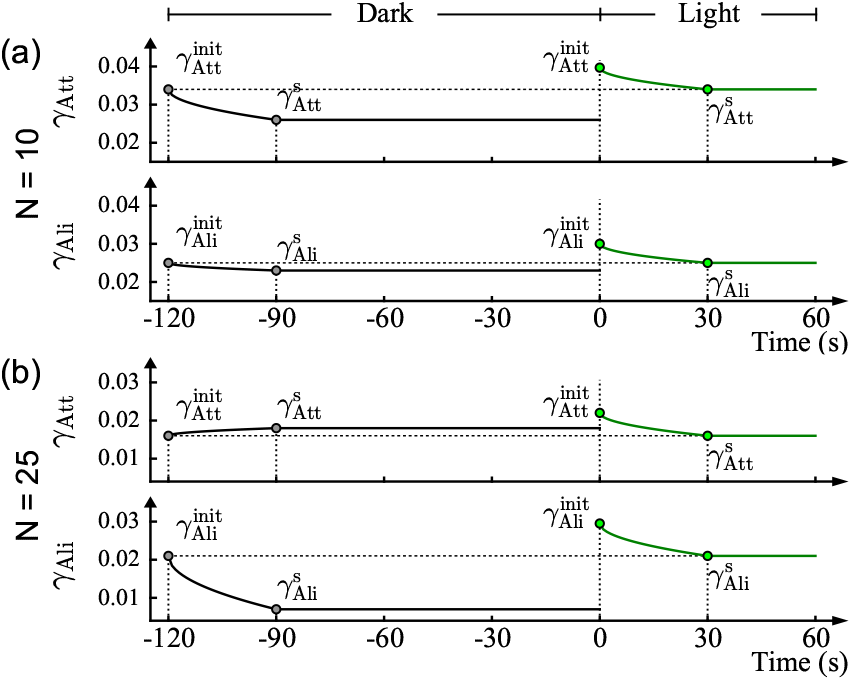
Time profiles of the strengths of attraction *γ*_Att_ and alignment *γ*_Ali_ interactions used in the numerical simulations for (a) *N* = 10 and (b) *N* = 25. After each increase or decrease in light, the intensities of *γ*_Att_ and *γ*_Ali_ vary from their initial values during a habituation period of 30 seconds and then keep a constant value. The initial parameter values in the Dark period are exactly the same as those at the end of the previous Light period. When the light is switched on, the attraction and alignment strengths are assumed to take initial values 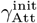 and 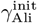 and then decrease to reach 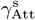 and 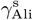 over a period of 30 seconds.

Figs. 2 and 3 reveal a good agreement between the experimental (solid lines) and simulation results (dotted lines) for all three conditions, including in the moments that immediately follow the changes in light intensity, when the fast collective response of fish to environmental changes is challenging to reproduce (see also SM Movie S5, S6, S7 and S8 [25]). In particular, the model reproduces the PDF of polarization, milling, and dispersion for all lighting conditions and for both group sizes. Figs. 8 and 9, also show that the model reproduces the distance of fish to the wall ⟨*r*_w_ ⟩*/R* and the distance of fish to the nearest neighbor (NND) observed in the experiments.

In the Stressing condition, the agreement is excellent in the 60 s before the light is turned on and in most of the cases even during the 2 min at 0.5 lx, as shown in Fig. 2. This is also the case during the 30 s following the transition time, *i*.*e*., in the last 30 s during which the light is at 25 lx. During the interval immediately following light changes, the model takes some time to adapt to the new light intensity (a short delay of 10 to 20 s depending on the observable) to adjust to the experimental value.

Together, these results suggest that the modulation of the collective behavior of fish in the Stressing condition results from the variation of amplitude of the spontaneous heading change of fish and the strength of the social interactions between fish.

### C. Impact of a mild stress on social interactions and phase diagram

The behavior of a dynamical system near phase transitions and the definition of critical states are better established in systems with a large number of particles, ideally in the infinite size limit. Although we are working with quite small systems, via the agent-based model, we demonstrate that our fish shoals tune towards critical points.

Figs. 5 and 6 show the phase diagrams for the polarization, milling, and dispersion, in the Dark and Light periods of the Stressing condition, and in the Control condition for each group size. They are represented in the *γ*_Att_-*γ*_Ali_-plane, together with the diagrams of fluctuations of each quantity, *χ*_*P*_ and *χ*_*M*_ (in physics, assimilated as susceptibilities), measured by the variance of the corresponding order parameter times the group size. The value of *γ*_R_ is also indicated for each condition. In each diagram, the value 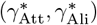 that better reproduces the experimental results in our simulations is represented by a *white cross* symbol.

**FIG. 5.**
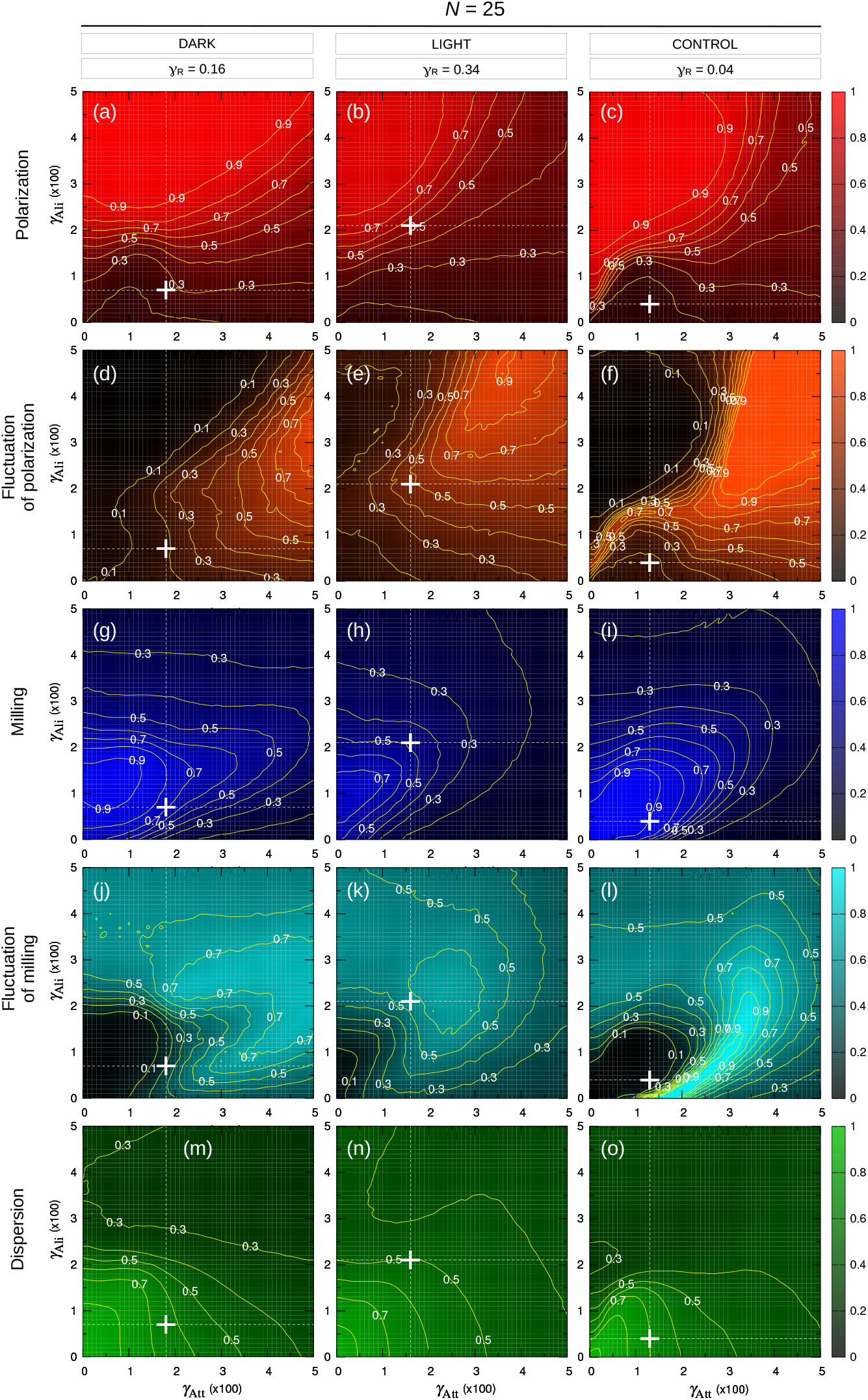
Polarization (a-c), fluctuations of polarization (d-f), milling (g-i), fluctuations of milling (j-l), and dispersion (m-o) as functions of the intensity of attraction *γ*_Att_ and alignment *γ*_Ali_, for the Dark (left) and Light (middle) periods of the Stressing condition and the Control condition in groups of *N* = 25 fish. Fish interact with their two most influential neighbors. White cross and dashed white lines indicate the value 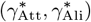 that best reproduces the experimental results in the numerical simulations. The value of the noise *γ*_R_ for each condition is also shown. The estimated interaction parameters are far from the critical region for the Dark and Control conditions, and lie in this region is the Light condition.

**FIG. 6.**
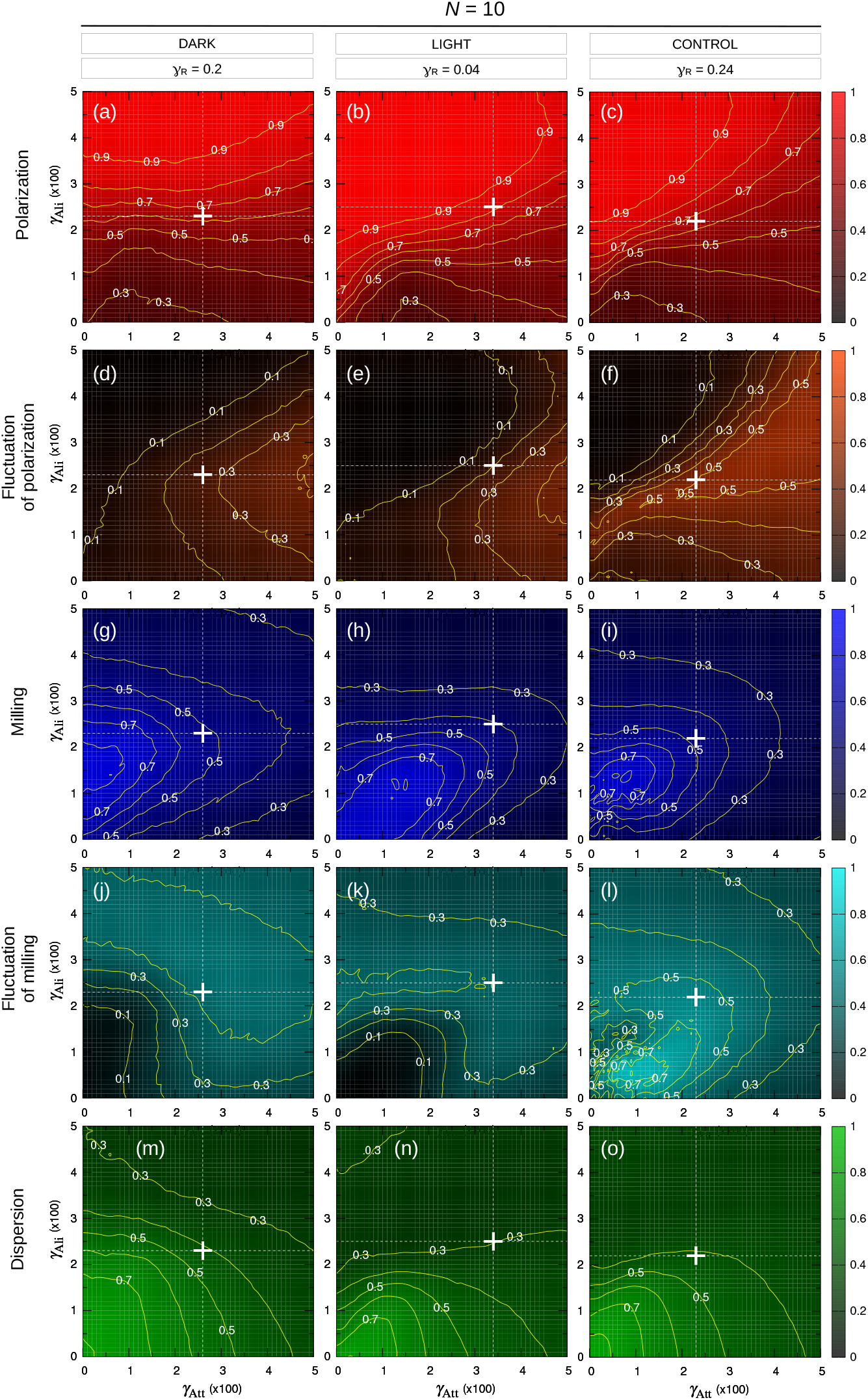
Polarization (a-c), fluctuations of polarization (d-f), milling (g-i), fluctuations of milling (j-l) and dispersion (m-o) as functions of the intensity of attraction *γ*_Att_ and alignment *γ*_Ali_, for the Dark (left) and Light (middle) periods of the Stressing condition and the Control condition in groups of *N* = 10 fish. Fish interact with their two most influential neighbors. White cross and dashed white lines indicate the value 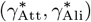 that best reproduces the experimental results in the numerical simulations. The value of the noise *γ*_R_ for each condition is also shown. In the three conditions, the estimated interaction parameters lie in the critical region and the 3 phase diagrams are relatively similar, contrary to what is found for *N* = 25.

Let us first consider the experiments performed with *N* = 25 fish. Figs. 5(a) and (g) show that in the Dark period of the Stressing condition, the system is in a state where the polarization is low (*P <* 0.3) and the milling is relatively high (*M >* 0.7). What is important here is not the absolute values of *P* and *M*, but their relative position with respect to the transition region between phases, *i*.*e*., the location of the white cross with respect to where the contour lines are more tightly packed. In the polarization diagram [Fig. 5(a)], the system is in the dark-red region, well below the transition line, and in the milling diagram [Fig. 5(g)], the system is almost at the top region in light-blue. This situation corresponds to a relaxed state (*i*.*e*., non-stressed), almost identical to the one observed in the Control condition [Fig. 5(c, i), *P <* 0.2, *M* = 0.9], which represents the system’s relaxed state when no external stress is applied.

When the light is switched on (*i*.*e*., in the Light period of the Stressing condition), the system jumps into the transition region, where small changes in the parameters induce significant changes in *P* and *M*. Comparing the three figures for each light condition and in both measures *P* [Fig. 5(a, b, c)] and *M* [Fig. 5(g, h, i)], the white cross clearly shifts from the interior of a phase into the transition region. In the diagrams of fluctuations [Fig. 5(d, e, f) and Fig. 5(j, k, l)], the white cross is located in the region of small values in the Dark and Control conditions, while in a region of higher values in the Light condition (not in the region of highest fluctuations, due to the limited size of the system). While this jump may seem small here, it is quite significant for such a small system.

This scenario corresponds to what one would expect from a biological perspective. Initially, the fish are in a non-stressed resting state. When the stressing stimulus is applied, they move to a critical state in which they can quickly adjust their behavior. Then, after the stimulus, they progressively relax, eventually returning to the Control state.

When a system is close to a transition region, the relaxation times tend to be longer (ultimately diverging in very large systems). Despite the small size of our system, this behavior is clear in the representation of the dynamics shown in Fig. 2(d, e, f). When suddenly changing from the Dark to the Light period in the Stressing condition, all three measures, polarization, milling, and dispersion, abruptly shift to new values, and then slowly relax towards the values of the Control condition. This gradual relaxation is particularly noticeable in the dispersion shown in Fig. 2(f). Here, the relaxation interval is too short to reach the stationary state. In turn, when the system moves back from Light to Dark, the relaxation time is much shorter, as expected when far from a transition region. We estimated these relaxation times (see SM [25], Fig. S2), showing that the relaxation from the critical state in Light is approximately one order of magnitude longer than when returning to Dark (*τ* = 95.3 s *vs. τ* = 12.7 s), thus suggesting that in the Light condition the system is indeed close to a phase transition.

For *N* = 25, the situation is thus as follows: the fish are initially calm, then when a stress stimulus occurs, they quickly shift closer to the transition region to be able to quickly react to potential danger. If nothing happens, they gradually relax, slowly returning to their steady state.

The situation is different for *N* = 10. Fig. 6 shows that the changes in polarization and milling are smaller than in the larger group. In fact, the three phase diagrams, corresponding to each light condition, are nearly identical in terms of both *P* [Fig. 6(a, b, c)] and *M* [Fig. 6(g, h, i)]. In addition, the fluctuation diagrams for the Dark and Light periods in the Stressing condition are also similar [Fig. 6(d, e) and Fig. 6(j, k)]. Although a slight increase in *P* is observed in the Light period, the interaction parameters (white cross) remain essentially in the same place, inside the transition region. Thus, in the three phase diagrams, the parameters corresponding to the experiment are located in the transition regions for the polarization and milling order parameters. In fact, since the parameters for the Dark and Control conditions are already in the transition region, it is not surprising to also find that the parameters for the stressed Light condition lie in the same regions. This suggests that in such small groups, the fish are already experiencing a stress state in both the Dark period and the Control condition, and that switching on the light only induces a small additional stress. One possible explanation is that groups of only 10 individuals experience an inherently stressful condition because the typical group size for this fish species in nature is around 30 individuals. It could also be that, since a group of 10 has a higher proportion of individuals near the periphery compared to a group of 25, a certain level of anxiety may persist. A social buffering effect, where the presence of others reduces stress levels in social species, could explain why smaller groups experience more stress [23, 24, 29–31]. Thus, the reduced behavioral changes across the different light conditions (all in the transition regimes) may reflect a state of constant vigilance rather than a true transition in the response to an additional stress.

## III. DISCUSSION

Identifying the mechanisms by which animal groups change their collective state is a key step to understanding how these systems adapt to environmental conditions [32]. In this work, we have investigated the mechanisms by which fish schools tune their collective state in response to controlled mild stress. Previous works have found signatures of critical state in fish schools through the analysis of spontaneous behavioral cascades, where large turns in the direction of motion of individuals propagate across a group [19, 20]. The power-law distributions of cascade duration, size and inter-event times have provided evidence of scale-free behavior in schooling fish, suggesting that these systems were operating near a phase transition between two collective states. In this region of the parameter space, the school exhibits multistability and regularly shifts between the two states, its responsiveness to perturbations is maximum and information can quickly propagate across arbitrarily large scales [9–11]. Other works have pointed out that fish schools can also regulate the distance to criticality according to the riskiness and noisiness of the environment [21].

Here, we used a data-driven computational modeling approach to quantify where in the parameter space groups of rummy-nose tetras operate both in an unperturbed condition and when all fish experience stress. We also studied the impact of group size on individual and collective behavior. In each condition, we measured the polarization, milling, and cohesion of the group. We then used the burst-and-coast model introduced in [27, 33, 34] to account for collective swimming and light-induced modulation of social interactions [28] in *H. rhodostomus* to determine the values of noise (*γ*_R_), attraction (*γ*_Att_) and alignment (*γ*_Ali_) parameters that lead to the experimentally measured values of polarization, milling, and cohesion under these conditions.

Our experimental and simulation results indicate that the collective state of fish and their reaction to a stressing condition depend on the size of the group to which they belong. In the Control condition and also in the Dark period of the Stressing condition, while groups of 25 fish are clearly in the milling phase (*i*.*e*., far from a critical region), groups of 10 fish are located in the transition region for the polarization and milling order parameters suggesting that individuals are experiencing a stress state presumably because they usually swim in larger groups in the wild [35] or in the fish facility. These results suggest that group size, presumably through a social buffering effect, modulates the strength of interactions between fish and their level of stress [23, 29, 30]. Empirical studies have shown that the presence of peers is effective in reducing an individual’s cortisol or corticosterone response to stress [24, 31], thus making individuals living in groups more relaxed/less stressed than when they are alone. A similar group size effect has been demonstrated in other species such as *Kuhlia mugil* in which an increase in fish density within a tank leads to a decrease in the strength of social interactions [36].

Moreover, when the transition between the Dark and the Light period of the Stressing condition occurs, in groups of 10 fish individuals react by increasing their randomness (*i*.*e*., increasing the amplitude of the spontaneous heading change) and their tendency to move closer to their peers, while in groups of 25 fish, individuals decrease their randomness and strengthen their tendency to stay aligned with their peers. However, while groups of 10 fish remain inside the transition region between the two phases of schooling and milling, groups of 25 fish move from the milling phase into the transition region when light is switched on. It is precisely in this zone of the parameter space that the fluctuations of the schooling and milling parameters are the largest [Fig. 5(d, e, f), Fig. 5(j, k, l)], and where the sensitivity and reactivity of the shoal are maximal [9–11]. Altogether, these results suggest that when fish are stressed, they readily tune the strength of their social interactions so that the group collectively reaches a critical state which allows a fast adaptation of its behavior in response to changes occurring in the environment. However, as the level of stress of fish is itself modulated by the size of the group, a small group of 10 individuals will immediately operate in a critical state while an additional level of stress induced by the sudden change of illumination is needed to lead a larger group of 25 fish in the critical regime. Furthermore, a large group will allow individuals to quickly return to a relaxed state, whereas in small groups, fish are in a relatively higher level of stress, which will maintain the group in a critical state.

Our data-driven burst-and-coast model accurately reproduces not only qualitatively but also quantitatively the state transition of fish under stress in the two group sizes, including the temporal changes and probability density distributions of several fine observables used to characterize the collective state of the fish.

Some work have suggested that one possible mechanism used by fish to regulate their collective state under perceived risk relies on a change in the structure of the interaction network between fish [21, 37]. Our findings show that the stress of fish associated with a perceived risk, but also the size of the group modulate the intensity of social interactions without the fish having to change the structure of the interaction network which remains fixed regardless of the environment, with each fish only interacting with its two most influential neighbors. This constitutes an alternative mechanism for the regulation of collective state in fish schools, which is also much simpler from a cognitive perspective because it keeps the cognitive load of fish constant [33, 38].

In conclusion, this work made it possible to reveal and model the mechanisms underlying collective state transition phenomena observed in fish [39–41] and the conditions under which these animal groups can reach a critical state.

## IV. MATERIALS AND METHODS

### A. Experiments and data collection

a. *Ethics statement*. Experiments were approved by the local ethical committee for experimental animals were performed in an approved fish facility (A3155501) under permit APAFIS#27303-2020090219529069 v8 in agreement with the French legislation.
b. *Study species*. Rummy-nose tetras (*Hemigrammus rhodostomus*) were purchased from Amazonie Labège in Toulouse, France. This is a small freshwater fish that forms highly cohesive and polarized schools [42, 43]. Fish were kept in 16 L aquariums on a 12:12 hour, dark:light photoperiod, at 24.9^*°*^C (± 0.8^*°*^C) and were fed *ad libitum* with fish flakes. The average body length (BL) of the fish used in these experiments is 3.1 cm.
c. *Experimental setup*. We used a rectangular glass tank of 120 cm × 120 cm supported by a structure of metal beams 20 cm high, inside which was installed a circular white arena of radius *R* = 25 cm. The tank was filled with 7 cm of water of controlled quality (50% of water purified by reverse osmosis and 50% of water treated by activated carbon) heated at 27.11 ^*°*^C (± 0.54 ^*°*^C). The tank was surrounded by white opaque curtains and the experimental room was lit by portable light sources, ensuring uniform lighting. The illumination of the circular arena was remotely controlled to produce either a constant level of light (25 lx) or an intermittent lighting (0.5 lx, 25 lx). Fish movements were recorded with a Sony HandyCam HD camera from above the setup at 30 Hz (1500*×*1500 p) in the control experiments and at 25 Hz (1920*×*1080 p) in the intermittent light experiments.
d. *Experimental design*. For each group size *N* = 10 and 25 fish, we carried out a total of 10 experiments that included 4 replicates in the *Control* condition, where light intensity was kept constant at 25 lx for one hour, and 6 replicates in the *Stressing* condition, in which light intensity alternates over ten periods of 3-minutes consisting of two minutes at low light intensity (0.5 lx) followed by one minute at a high light intensity (25 lx). The transition between light intensities was instantaneous and remotely controlled (see Fig. 1). The duration of the low light intensity time interval was chosen so that fish can quickly recover from the Stressing condition and adopt their default state when they are in the dark. The duration of the high light intensity interval was chosen so that fish had time to react to the change in brightness and adapt their behavior to cope with the Stressing condition. The number of periods in the stress conditions was large enough to allow the acquisition of experimental data within a reasonable time, and small enough to avoid possible conditioning or habituation effects to intermittent lighting; see the boxplots in SM Fig. S1 [25]. Before each session, fish were introduced in the experimental tank with an acclimation of 10 min at 25 lx in the Control condition and at 0.5 lx in the Stressing condition. In all experimental runs performed on the same day, we used different fish, and the intermittent light experiments were spaced out by one week to prevent conditioning or habituation phenomena.
e. *Data extraction and pre-processing*. Raw data thus represent a total of 14 h of video. We then used FastTrack [44] to extract fish trajectories from the videos (*i*.*e*., the 2D position of each fish at each frame). Due to the frequent cross-occlusion between individuals and the misidentification of individuals in low light condition, the tracking accuracy (*i*.*e*., the percentage of frames in which all *N* fish have been correctly identified) varies between 90% and 99% in the Control condition, and 51% and 91% in the Stressing condition (see SM Table S1 [25] for details on tracking accuracy and [28] in which the same tracking procedures were used).

### B. Quantification of collective behavior

We characterize the collective behavioral patterns exhibited by fish groups by means of three observables.

We first define 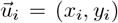 and 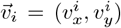 as the position and velocity vectors of fish *i* [see Fig. 7(a)], and 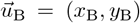 and 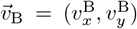 as the position and velocity vectors of the barycenter B of the group, where

**FIG. 7.**
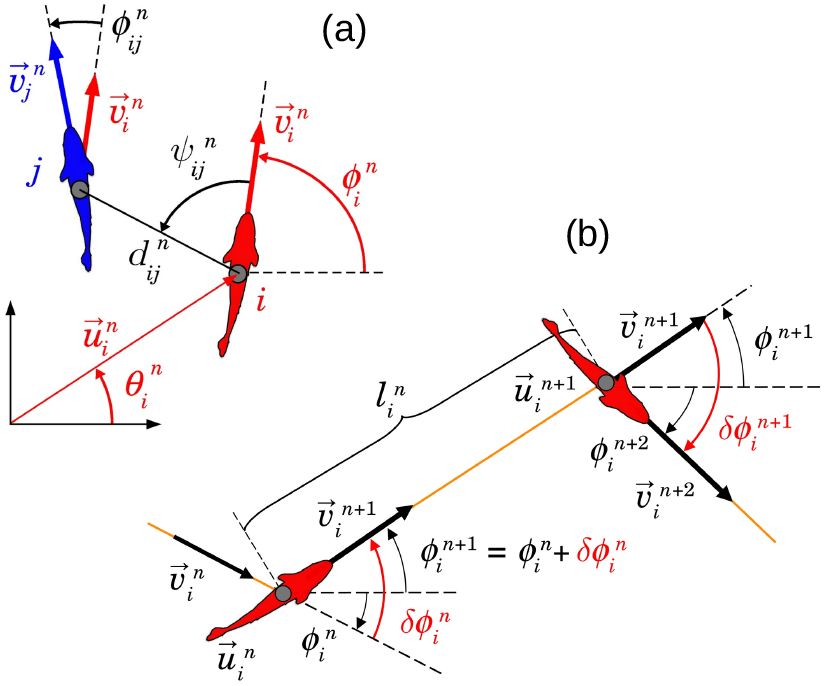
Variables used to describe social interactions between fish and their burst-and-coast swimming mode. (a) Individual (red) and social (black) state variables of fish *i* with respect to fish *j* at the instant 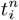 when fish *i* performs its *n*-th kick: 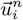 and 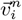 are the position and velocity vectors of fish *i*, 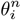 and 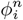 are the angles that these vectors make with the horizontal line,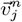 is the velocity vector of fish *j* at time 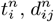 is the distance between fish *i* and fish *j* 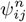 is the angle with which fish *j* is perceived by fish *i* (not necessarily equal to 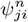), and 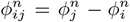 is the heading difference between both fish. (b) Schematic of the *n*-th kick performed by fish *i* moving from 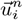 at time 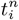to 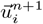 at time 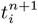along a distance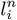. Orange lines denote fish trajectory, black wide arrows denote velocity vector, curved arrows represent angles. The heading angle change of fish *i* at time *t* is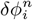. Fish heading during its *n*-th kick is 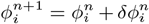. Red angles show the heading variation of the fish at the kicking instants 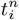 and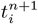.

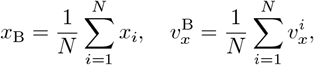

with similar expressions for *y*_B_ and 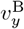. We assume that the heading angle of fish *i* is given by its velocity vector, *ϕ* = `ATAN2` (*v*_*y*_, *v*_*x*_). Similarly, the heading angle the barycenter of the group is given by 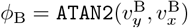

The three dimensionless observables used to quantify the group behavior at instant *t* are defined as follows:

- Dispersion *D*(*t*):

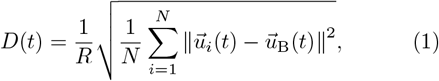

where *R* is the radius of the fish tank. Large values of *D*(*t*) mean that fish are dispersed in the tank, while small values mean that they are aggregated.
- Polarization *P*(*t*) ∈ [0, 1]:

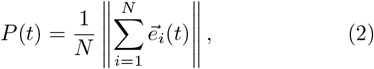

where 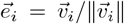 is the unit vector in the direction of motion of fish *i*. Values of *P*(*t*) close to 1 mean that *N* fish are aligned and move in the same direction. Values of *P* close to 0 indicate that the *N* vectors point in different directions, but can also mean that vectors are collinear and in opposite directions so that they cancel each other (*e*.*g*., when half of the vectors pointing north and the other half pointing south, like when fish are milling). Configurations in which fish are randomly aligned give rise to values of *P* of order 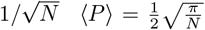 and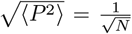 for truly uncorrelated headings).
- Milling *M*(*t*) ∈ [0, 1]:

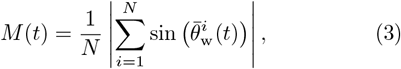

where 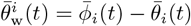. Variables with a bar are defined in the barycenter system of reference: 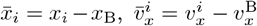 and similar expressions are used for the *y*-components. The relative position and heading angles of fish *i* with respect to B are respectively 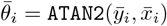 and 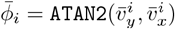. The milling *M*(*t*) measures how much the fish turn in the same direction around the center of the tank, independently of the direction of rotation. In order to estimate the sensitivity of fish to light stimuli, we measure the variance of these observables over a period of time, times the group size,

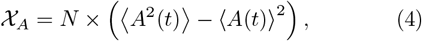

where symbol ⟨·⟩ means average over time and *A* can be the dispersion *D*, the polarization *P*, or the milling *M*.

We also used two additional observables, the distance of each fish to the wall 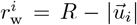, where *R* is the radius of the tank, and the distance of fish to the nearest neighbor NND. Figs. 8 and 9 show respectively the time series and PDF of these observables.

**FIG. 8.**
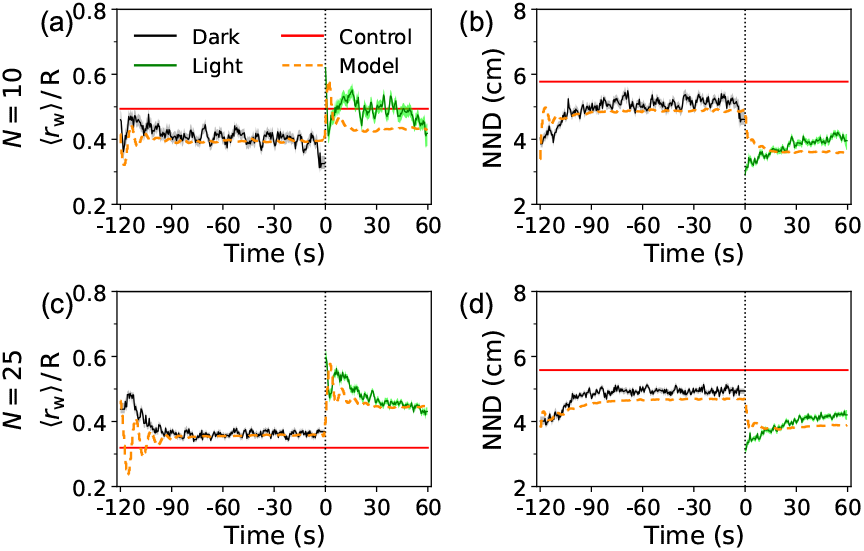
Time series of the average distance of fish to the wall ⟨*r*_w_⟩ */R* and the average distance of fish to the nearest neighbor (NND) in the experiments. (a,b) *N* = 10 fish and (c,d) *N* = 25 fish in the Dark (black lines) and Light (green lines) periods of the Stressing condition and in the Control condition (red lines). Shadows represent the 68% confidence interval (bootstrap of 1000 rounds) for experimental results. Orange dashed lines represent the model simulations.

**FIG. 9.**
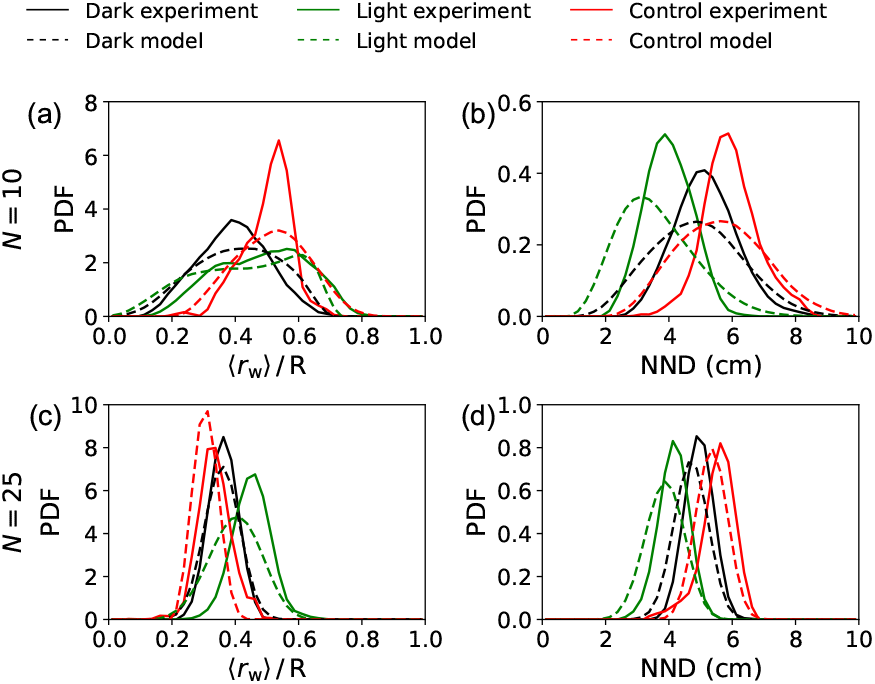
Probability density functions (PDF) of the distance to the wall ⟨*r*_w_⟩ */R* and the distance to the nearest neighbor (NND) observed in the experiments (solid lines) and obtained in the numerical simulations (dashed lines). (a,b) *N* = 10 fish and (c,d) *N* = 25 fish in the Dark (black lines) and Light (green lines) periods of the Stressing condition and in the Control condition (red lines).

### C. Computational model

We use the burst-and-coast model introduced in [27] and [33, 34] to account for collective swimming behavior in *Hemigrammus rhodostomus* and extended in [28] to account for the impact of varying levels of light intensity on social interactions between fish.

As found in [28], the model can remarkably reproduce the experimental results of fish swimming under different light intensities (0.5, 1, 1.5, 5, and 50 lx) for a wide range of group sizes (*N* = 1, 2, 5, and 25). Both the structure of the model and the shape of the interaction functions (*i*.*e*., their analytical expressions) are preserved here, the simple adjustment of the strength and range parameters of the interactions being sufficient to reach an excellent agreement between the numerical simulations and the experimental results.

The model, reflecting the burst-and-coast swimming mode of rummy-nose tetra, considers that fish trajectories are made of a succession of segments, along which fish keep their orientation while decelerating due to the water drag. The short events between segments (of typical duration 0.1 s for real fish; considered as instantaneous in the model) during which a fish changes both its velocity and also its direction of motion are called “kicks”. The individual state of a fish *i* at the instant 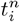 when it performs its *n*-th kick is determined by its position and its heading angle,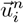 and 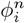, respectively. Precisely at time 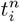, the fish chooses its new heading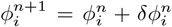, where 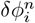 is the heading angle change, and glides along a straight line of length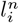, with an initial speed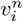, and during a time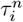.

At the end of the kick, at time 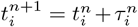, the new position and heading are given by :

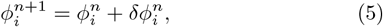

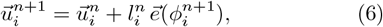

where 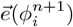 is the unit vector of the direction of motion of the fish during its *n*-th kick, given by 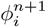[see Fig. 7(b)].

In this model, the length, initial speed, and the duration of the kicks performed by a fish are independent of those performed by other fish. The duration and speed are sampled from bell-shaped distributions resembling the experimental ones, and given by

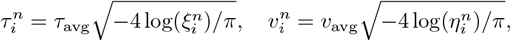

where τ_avg_ and *l*_avg_ are the average kick duration and length measured in the experiments, and 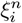 and 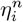 are two random numbers uniformly distributed in (0, 1). Assuming an exponential decay of the speed between kicks (fully consistent with experiment [27]), the peak speed 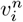 and the kick duration and length 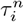 and 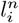 are linked by the relation, *l* = *vτ*_0_[1 − exp(−*τ/τ*_0_)], where *τ*_0_ is a decay time. Then, the position of fish *i* at any intermediate time 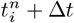 in 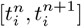 can be calculated as follows,

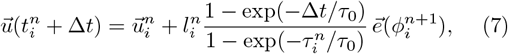

where 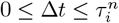.

The heading angle change of an individual fish *ϕ* results from the additive combination of its spontaneous behavior, the physical constraints of the environment (*e*.*g*., obstacles), and the social interactions with other fish:

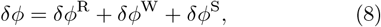

where indices R, W, and S stand for Random, Wall, and Social, respectively. The analytical expressions of these forces were derived in [27] for pairwise interactions by means of a procedure of reconstruction applied to experimental data of pairs of fish. Here we use a simplified version introduced in [33] for groups of larger size.

The spontaneous heading change of fish can be described by a Gaussian noise when the fish swims alone far from the wall, and their amplitude is smaller when the fish comes close to the wall. Thus,

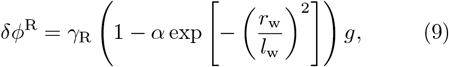

where *γ*_R_ is the amplitude of these heading changes, *l*_w_ is the distance of fish to the wall from which the amplitude is reduced, *α* is the strength of this reduction, and *g* is a Gaussian random variable with zero average and unit variance.

The wall exerts on the fish a repulsive force that depends only on the distance of the fish to the wall *r*_w_ and its relative orientation to the wall *θ*_w_. We assume that this dependence is decoupled, so that the heading angle change due to this force can be written as

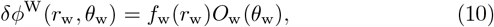

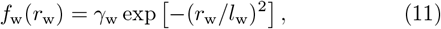

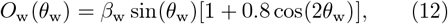

where *γ*_w_ is the intensity and *l*_w_ the range of the wall repulsion, and *β*_w_ is a constant of normalization of the angular function so that the mean of the squared function in [−*π, π*] is equal to 1: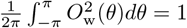.

The social interactions between fish *i* are considered to be the linear combination of pairwise interactions 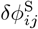 with a specific number of neighbors *j* = 1, …, *k*. The pairwise interactions 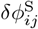 depend only on the relative state *s*_*ij*_ = (*d*_*ij*_, *ψ*_*ij*_, *ϕ*_*ij*_) of fish *j* with respect to fish *i*, given by the distance between them *d*_*ij*_, the viewing angle *ψ*_*ij*_ with which fish *i* perceives fish *j*, and their relative alignment *ϕ*_*ij*_ = *ϕ*_*j*_ *™ ϕ*_*i*_. It has been shown in [27, 33] that in *H. rhodostomus*, each fish interacts with its neighbors by means of an attraction interaction (negative/repulsive at short range) and an alignment interaction, and selectively interacts with the *k* = 2 specific neighbors having the largest and the second-largest associated value of 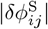. Thus,

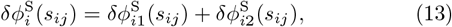

where

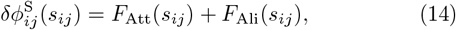

and

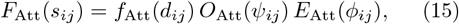

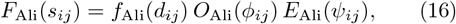

for *j* = 1, 2. The analytical expressions found for these functions were extracted from specific experiments under different uniform light intensities and are as follows [28]:

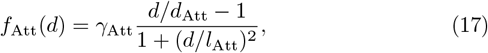

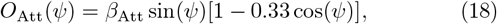

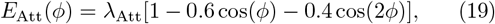

for the attraction, and, for the alignment:

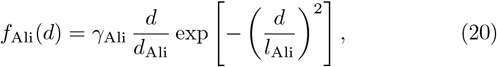

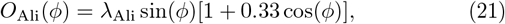

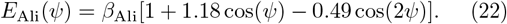

Here, *γ*_Att_ and *γ*_Ali_ are the dimensionless intensities of the attraction and alignment forces respectively, *d*_Att_ and *d*_Ali_ are the distance below which the corresponding interaction changes sign, *l*_Att_ and *l*_Ali_ are the respective ranges of action, and *β*_Att_, *λ*_Att_, *β*_Ali_ and *λ*_Ali_ are the constants of normalization of the corresponding angular function. The parameter values are given in Table II and Fig. 10.

**TABLE 2.** Model parameters used in the simulations of different illumination conditions and different group sizes.

| Parameter | $N = 10$ | | | $N = 25$ | | |
| --- | --- | --- | --- | --- | --- | --- |
|  | Dark | Light | Ctrl | Dark | Light | Ctrl |
| $\gamma_w$ | 0.3 | 0.4 | 0.4 | 0.3 | 0.4 | 0.4 |
| $\beta_w$ | 1.9612 | | | | | |
| $l_w$ (m) | 0.031 | 0.053 | 0.053 | 0.031 | 0.053 | 0.053 |
| $\alpha$ | 0.35 | 0.66 | 0.66 | 0.35 | 0.66 | 0.66 |
| $d_{\text{Att}}$ (m) | 0.015 | | | | | |
| $\beta_{\text{Att}}$ | 1.43 | | | | | |
| $\lambda_{\text{Att}}$ | 0.9 | | | | | |
| $l_{\text{Att}}$ (m) | 0.1 | 0.165 | 0.165 | 0.1 | 0.165 | 0.165 |
| $d_{\text{Ali}}$ (m) | 0.015 | | | | | |
| $\beta_{\text{Ali}}$ | 0.9 | | | | | |
| $\lambda_{\text{Ali}}$ | 1.43 | | | | | |
| $l_{\text{Ali}}$ (m) | 0.092 | 0.135 | 0.135 | 0.092 | 0.135 | 0.135 |
| $\tau_{\text{avg}}$ (s) | 0.32 | 0.55 | 0.55 | 0.32 | 0.55 | 0.55 |
| $\tau_0$ (s) | 0.34 | 0.86 | 0.86 | 0.34 | 0.86 | 0.86 |
| $l_{\text{avg}}$ (m) | 0.028 | 0.053 | 0.053 | 0.028 | 0.053 | 0.053 |

**FIG. 10.**
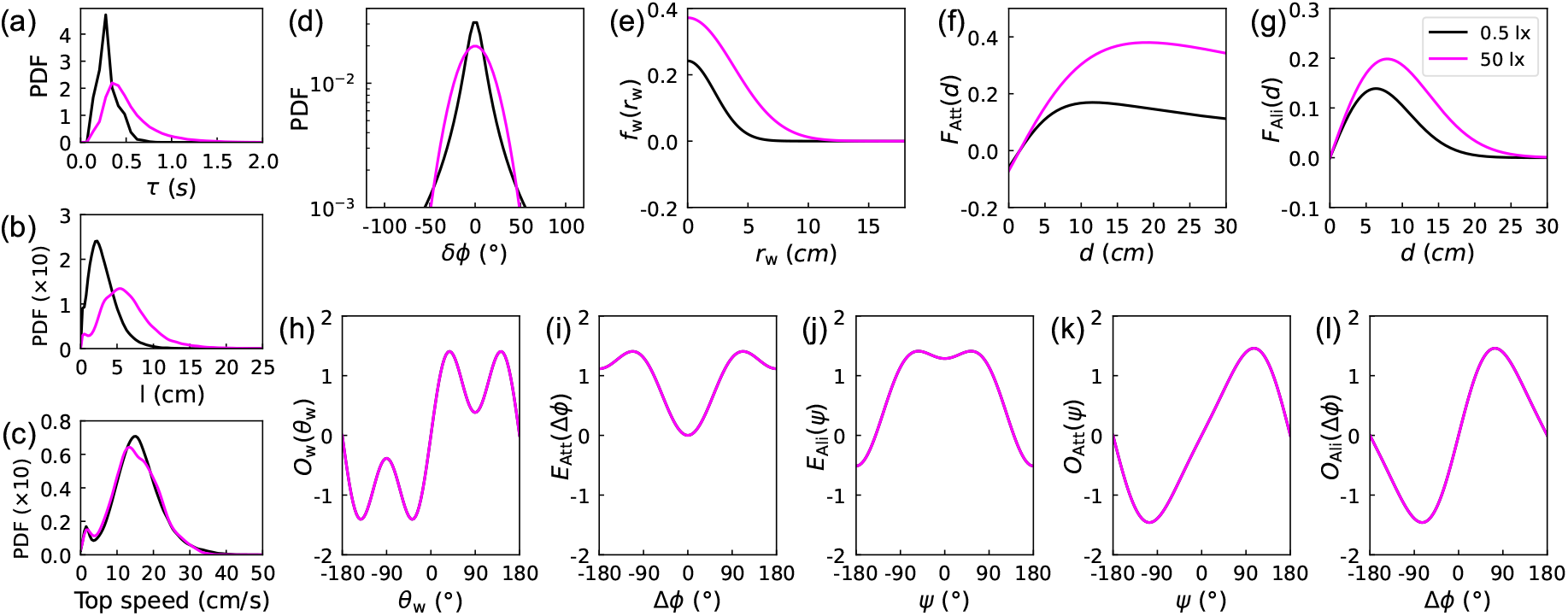
Functions used in the model to account for the effects of light intensity on the burst-and-coast swimming and the social interactions between pairs of fish derived from [28]. Probability density function (PDF) of (a) the kick duration *τ*, (b) the kick length *l*, (c) the peak speed at the kicking instant *v*, and (d) the spontaneous heading change *ϕ*. (e) Repulsion from the wall *f*_w_ (*r*_w_) as a function of the distance to the wall *r*_w_. (f,g) Attraction *F*_Att_(*d*) and alignment *F*_Ali_(*d*) as function of the distance between fish *d*. (h) Repulsion from the wall *O*_w_(*θ*_w_) as a function of the relative orientation of the fish to the wall *θ*_w_, (i,j) Attraction between fish *E*_Att_(Δ*ϕ*) and *O*_Att_(*ψ*) as a function of the relative orientation between fish Δ*ϕ* and the viewing angle *ψ* respectively, (k.l) Alignment between fish *O*_Ali_(*ψ*) and *E*_Att_(Δ*ϕ*) as functions of the viewing angle *ψ* and as a function of the relative orientation between fish Δ*ϕ. O* and *E* indicate the parity of the function, odd and even, respectively. Black lines correspond to 0.5 lx, pink lines to 50 lx.

Finally, a rejection procedure is used when the position of the fish calculated at the end of the kick is out of the tank. In that case, the kick is rejected, a new kick is calculated, a new angle of spontaneous variation *ϕ*^R^, a new kick length 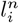 and a new kick duration 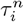 are sampled, until the final position at the end of the kick is inside the tank. if no suitable kick is found after 1000 tries, *ϕ*^R^ is randomly sampled from a uniform distribution (see [27] for more details).

### D. Numerical simulations and parameters estimation from experiments

Three series of numerical simulations of the burst-and-coast model were carried out. The first one was performed to get accurate values of the parameters (*γ*_R_, *γ*_Att_, *γ*_Ali_) for each condition: the Control condition (25 lx), the Dark period of the Stressing condition (0.5 lx) and the Light period of the Stressing condition (25 lx). In the second series of simulations, we run the model under each experimental condition, and the third series was performed to build the corresponding phase planes.

a. *Parameter values*. The parameter values used in the model are those determined in [28] for different light intensities and group sizes. Parameter values corresponding to the light intensity of 25 lx were obtained by interpolation from measurements made at 5 lx and 50 lx by [28]. Likewise, the parameter values for the groups of *N* = 10 were obtained by interpolation from measurements done in groups of 5 and 25 fish at different light intensities in [28]. Note, however, that in [28] the parameter values were measured with a constant illumination. Therefore, we have to take into account the modulation of individual parameters caused by sudden changes in light intensity. According to the results reported in [34], the amplitude of individuals’ spontaneous heading change, *γ*_R_, as well as the strength of social interactions between fish, *γ*_Att_ and *γ*_Ali_, can deeply affect the group behavior. Therefore, we assume that intermittent illumination causes changes in ***γ*** = (*γ*_R_, *γ*_Att_, *γ*_Ali_) without affecting other parameters. The estimation procedure to find out their values is given below. We used the mean values and the PDFs of the three observables *P, M*, and *D* to quantify the agreement between the numerical simulations of the model and the experimental results. For each light condition and group size, we run a large number of numerical simulations to find the triplet 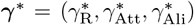 that minimizes the total error given by:

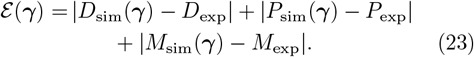 This is done in two steps. First, for each fixed value of *γ*_R_, we sweep through the plane *γ*_Att_-*γ*_Ali_ with a fine discretization and find the pair (*γ*_Att_, *γ*_Ali_) that yields the smallest error *ℰ* (*γ*_R_). Fig. 11 shows the resulting curves ℰ (*γ*_R_) for each group size and each light condition. Then, we sweep through the values of *γ*_R_ and the corresponding pairs (*γ*_Att_, *γ*_Ali_) obtained for each one and find the value of *γ*_R_ that yields the smallest error ℰ (***γ***^∗^). These values are represented by vertical dashed lines in Fig. 11, for each condition and each group size. We used a discretization of 10 values of *γ*_R_ with a step of 0.02 in different sub-intervals of [0, 0.42], depending on the group size and the condition, and a fine step Δ*γ* = 10^−3^ to sweep the *γ*_Att_-*γ*_Ali_ plane. The final parameter values are given in Table II The simulations carried out to find the value ***γ***^∗^ in the Control condition were performed in the same conditions as the experimental ones. We simulated 10 runs of 70 min at 25 lx, the first 10 min of each run corresponding to the habituation time. We only used the last 60 min to calculate the mean values *D*_sim_, *P*_sim_, and *M*_sim_. For the Stressing condition, numerical simulations were simplified with respect to what is done experimentally. Instead of simulating runs of 10 successive 3-minutes periods of 2 min in the Dark condition and 1 min in the Light condition, as in the experiments, we proceeded in two steps, first for the Dark condition, then for the Light condition. For the Dark condition, we simulated 600 simple runs of 2 min under constant light. For the Light condition, a jump in light must be considered in the 3-minutes period which is described below. In both Dark and Light conditions, each run started from an initial condition randomly selected from the initial states obtained in the Dark condition in the experiments. Then, the value of ***γ***^∗^ minimizing the error is extracted for each Dark and Light condition, using the respective simulated data. The simulations of the 3-minutes period in the Light condition were done as follows. According to [28], the three strength parameters *γ*_R_, *γ*_Att_, and *γ*_Ali_ take different values depending on the light intensity. A simple assumption could be made that, when transitioning from Dark to Light, these values jump from those corresponding to the initial light intensity to those corresponding to the new light intensity. Under this assumption, the variation of *γ*_Att_ and *γ*_Ali_ could be modeled using piecewise constant functions. However, the simulations of the model using this hypothesis show that the system is not able to adapt quickly enough to the change and remains in the stable state corresponding to the light intensity before the change. We thus considered that the values of *γ*_Att_ and *γ*_Ali_ undergo a larger jump and introduced, during the first 30 s of the Light period, a transition period during which *γ*_Att_ and *γ*_Ali_ smoothly decrease to the values corresponding to the final light intensity, allowing the fish group to progressively adapt its behavior to the changing values of *γ*_Att_ and *γ*_Ali_ (See Fig. 4). No transition is necessary for *γ*_R_. This mechanism used to account for the transition period was tested by comparing the results of the numerical simulations with the experimental ones. Both the duration of the transition period and the initial value have been determined after successive trials. We tried several decaying rates, and found that a square root behavior provided a smoother transition than a simple linear decay:

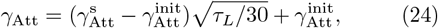

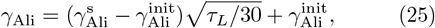

where *τ*_*L*_ ∈ [0, 30] is the time elapsed from the beginning of the Light condition.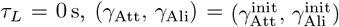, and at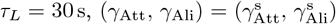. In the Light condition, 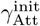 and 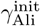 are set to 1.3 and 1.5 times the stable value, respectively. We run 600 simulations of one period composed of 2 min at 0.5 lx and 1 min at 25 lx, where the first 30 s of each interval corresponds to the transition from the initial state to the final one (Fig. 4). To calculate the mean values *P*_sim_, *M*_sim_, and *D*_sim_, we used the last 30 s of each interval of each condition, that is, the last 30 s of the 600 intervals of 2 min for the Dark condition, and the last 30 s of the 600 intervals of 1 min for the Light condition.
b. *Simulations of the experimental conditions*. Once the values of ***γ*** have been determined for each group size and illumination condition, a second series of numerical simulations was carried out to reproduce the experiments. These simulations were also used to find the adequate profile of the transition interval. For the Control condition, we ran 10 simulations of 70 min at 25 lx, the first 10 min being the habituation, and we used the last 60 min to calculate the mean values and the PDFs shown in Figs. 2 and 3. For the Stressing condition, we run 2400 simulations of one period of 3 min including the transition described above. The result is represented by the dashed lines in Figs. 2 and 3. To calculate the corresponding mean values and the PDFs shown in Fig. 3, we used the last 30 s of each interval of each light condition of the period, that is, the last 30 s of the 2400 intervals of 2 min for the 0.5 lx light intensity, and the last 30 s of the 2400 intervals of 1 min for the 25 lx light intensity. The agreement of the numerical simulations with the experimental results is shown in Figs. 2 and 3, for the three observables of polarization, milling and dispersion, and in Figs. 8 and 9 for the distance to the wall *r*_w_ and the distance to the nearest neighbor NND.
c. *Construction of the phase spaces*. Once the values of ***γ*** are known and the transition mechanism has been validated for each group size, we built the phase planes of each group size for the value of *γ*_R_ found with our iterative procedure. Mean values shown in the phase planes are calculated as in the numerical simulations. We performed 600 simulations of one period for each pair (*γ*_Att_, *γ*_Ali_), with a discretization step of Δ*γ* = 10^−3^ in [0, 0.05] × [0, 0.05], that is, 1.5 million runs for each phase plane and condition.

**FIG. 11.**
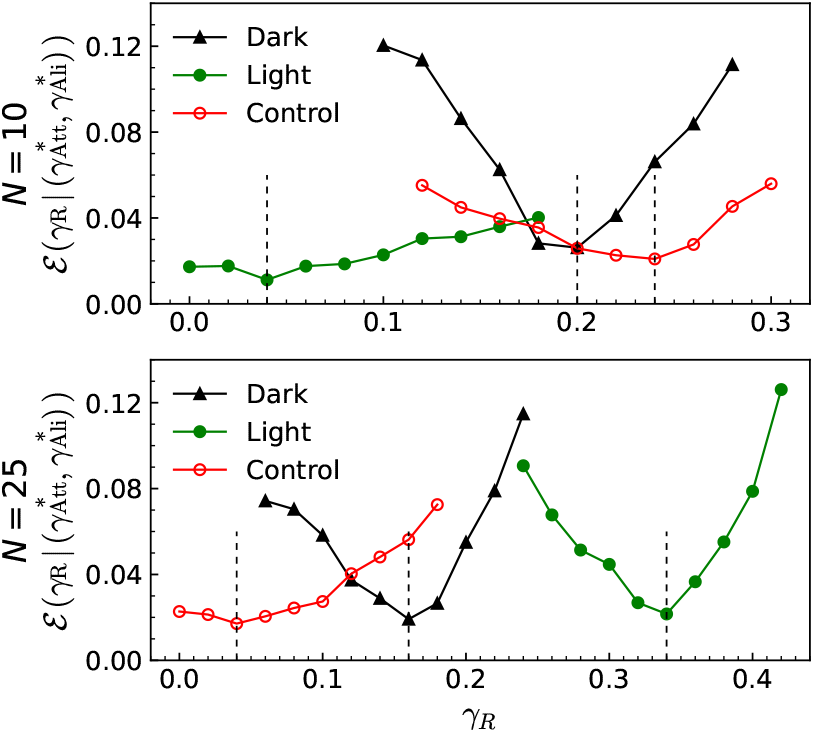
Determination of 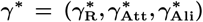, the triplet that minimizes the total error function ℰ defined in equation (23) as a function of *γ*_R_, for each condition and each group size. *γ*_Att_ and *γ*_Att_ have been optimized for each value of *γ*_R_ for *N* = 10 (top) and *N* = 25 (bottom) in the Dark (black lines) and Light (green lines) periods of the Stressing condition and in the Control condition (red lines). The vertical dotted lines indicate the value of *γ*_R_ that minimizes the error function.

## Supporting information

Supplementary Information for Experimental evidence of stress-induced critical state in schooling fish

## ACKNOWLEDGMENTS

This work was partly supported by the Indo-French Centre for the Promotion of Advanced Research (Project N°64T4-B) to GT and VG. G.L. was supported by a grant from the China Scholarship Council (CSC N° 202006040162). T.X. was supported by a grant from the China Scholarship Council (CSC N°202106040094). X.L. was supported by a grant from the China Scholarship Council (CSC N°202106040093). ZH was supported by the National Natural Science Foundation of China (Grant No. 62176022). GT, RE and CS were supported by the Agence Nationale de la Recherche (ANR-20-CE45-0006-1).

## Notes

### Competing Interest Statement

The authors have declared no competing interest.

## References

[1] Philip Ball. The self-made tapestry: pattern formation in nature. Oxford University Press, Inc., 1999.

[2] Mehdi Moussaid, Simon Garnier, Guy Theraulaz, and Dirk Helbing. Collective information processing and pattern formation in swarms, flocks, and crowds. Topics in Cognitive Science, 1(3):469–497, 2009.

[3] Scott Camazine, Jean-Louis Deneubourg, Guy Theraula, James Sneyd, and Nigel R Franks. Self-organization in biological systems. Princeton university press, 2020.

[4] Roland Wedlich-Söldner and Timo Betz. Self-organization: the fundament of cell biology. Philosophical Transactions of the Royal Society B: Biological Sciences, 373(1747):20170103, 2018.

[5] Eric Karsenti. Self-organization in cell biology: a brief history. Nature Reviews Molecular Cell Biology, 9(3):255–262, 2008.

[6] C Wolf and David Edmund Johannes Linden. Biological pathways to adaptability–interactions between genome, epigenome, nervous system and environment for adaptive behavior. Genes, Brain and Behavior, 11(1):3–28, 2012.

[7] H Eugene Stanley. Phase transitions and critical phenomena, volume 7. Clarendon Press, Oxford, 1971.

[8] Thierry Mora and William Bialek. Are biological systems poised at criticality? Journal of Statistical Physics, 144:268–302, 2011.

[9] Daniel S Calovi, Ugo Lopez, Paul Schuhmacher, Hugues Chaté, Clément Sire, and Guy Theraulaz. Collective response to perturbations in a data-driven fish school model. Journal of The Royal Society Interface, 12(104):20141362, 2015.

[10] Miguel A Munoz. Colloquium: Criticality and dynamical scaling in living systems. Reviews of Modern Physics, 90 (3):031001, 2018.

[11] Pascal P Klamser and Pawel Romanczuk. Collective predator evasion: Putting the criticality hypothesis to the test. PLoS Computational Biology, 17(3):e1008832, 2021.

[12] John M Beggs. The criticality hypothesis: how local cortical networks might optimize information processing. Philosophical Transactions of the Royal Society A: Mathematical, Physical and Engineering Sciences, 366(1864):329–343, 2008.

[13] John M Beggs and Nicholas Timme. Being critical of criticality in the brain. Frontiers in Physiology, 3:163, 2012.

[14] Enrique Balleza, Elena R Alvarez-Buylla, Alvaro Chaos, Stuart Kauffman, Ilya Shmulevich, and Maximino Aldana. Critical dynamics in genetic regulatory networks: examples from four kingdoms. PLoS One, 3(6):e2456, 2008.

[15] Bryan C Daniels, Hyunju Kim, Douglas Moore, Siyu Zhou, Harrison B Smith, Bradley Karas, Stuart A Kauffman, and Sara I Walker. Criticality distinguishes the ensemble of biological regulatory networks. Physical Review Letters, 121(13):138102, 2018.

[16] Giovanna De Palo, Darvin Yi, and Robert G Endres. A critical-like collective state leads to long-range cell communication in dictyostelium discoideum aggregation. PLoS Biology, 15(4):e1002602, 2017.

[17] William Bialek, Andrea Cavagna, Irene Giardina, Thierry Mora, Oliver Pohl, Edmondo Silvestri, Massimiliano Viale, and Aleksandra M Walczak. Social interactions dominate speed control in poising natural flocks near criticality. Proceedings of the National Academy of Sciences, 111(20):7212–7217, 2014.

[18] Alessandro Attanasi, Andrea Cavagna, Lorenzo Del Castello, Irene Giardina, Stefania Melillo, Leonardo Parisi, Oliver Pohl, Bruno Rossaro, Edward Shen, Edmondo Silvestri, et al. Finite-size scaling as a way to probe near-criticality in natural swarms. Physical Review Letters, 113(23):238102, 2014.

[19] Luis Gómez-Nava, Robert T Lange, Pascal P Klamser, Juliane Lukas, Lenin Arias-Rodriguez, David Bierbach, Jens Krause, Henning Sprekeler, and Pawel Romanczuk. Fish shoals resemble a stochastic excitable system driven by environmental perturbations. Nature Physics, 19:663– 669, 2023.

[20] Andreu Puy, Elisabet Gimeno, David March-Pons, M. Carmen Miguel, and Romualdo Pastor-Satorras. Signatures of criticality in turning avalanches of schooling fish. Physical Review Research, 6:033270, Sep 2024.

[21] Winnie Poel, Bryan C Daniels, Matthew MG Sosna, Colin R Twomey, Simon P Leblanc, Iain D Couzin, and Pawel Romanczuk. Subcritical escape waves in schooling fish. Science Advances, 8(25):eabm6385, 2022.

[22] Pawel Romanczuk and Bryan C Daniels. Phase transitions and criticality in the collective behavior of animals—self-organization and biological function. In Order, Disorder and Criticality: Advanced Problems of Phase Transition Theory, pages 179–208. World Scientific, 2023.

[23] Brett M Culbert, Kathleen M Gilmour, and Sigal Balshine. Social buffering of stress in a group-living fish. Proceedings of the Royal Society B, 286(1910):20191626, 2019.

[24] Kathleen M Gilmour and Brittany Bard. Social buffering of the stress response: insights from fishes. Biology Letters, 18(10):20220332, 2022.

[25] See supplemental material at [url will be inserted by publisher] for 2 supplementary figures, 2 supplementary tables, and 8 supplementary movies.

[26] C Aimon, Christophe Lebigre, Stephane Le Floch, and Guy Claireaux. Effects of dispersant-treated oil upon behavioural and metabolic parameters of the anti-predator response in juvenile european sea bass (dicentrarchus labrax). Science of The Total Environment, 834:155430, 2022.

[27] Daniel S Calovi, Alexandra Litchinko, Valentin Lecheval, Ugo Lopez, Alfonso Pérez Escudero, Hugues Chaté, Clément Sire, and Guy Theraulaz. Disentangling and modeling interactions in fish with burst-and-coast swimming reveal distinct alignment and attraction behaviors. PLoS Computational Biology, 14(1):e1005933, 2018.

[28] Tingting Xue, Xu Li, Guozheng Lin, Ramón Escobedo, Zhangang Han, Xiaosong Chen, Clément Sire, and Guy Theraulaz. Tuning social interactions’ strength drives collective response to light intensity in schooling fish. PLoS Computational Biology, 19(11):e1011636, 2023.

[29] Robert M Ross, Thomas WH Backman, and Karin E Limburg. Group-size-mediated metabolic rate reduction in american shad. Transactions of the American Fisheries Society, 121(3):385–390, 1992.

[30] Michael B Hennessy, Sylvia Kaiser, and Norbert Sachser. Social buffering of the stress response: diversity, mechanisms, and functions. Frontiers in Neuroendocrinology, 30(4):470–482, 2009.

[31] Natália Pagnussat, Angelo L Piato, Isabel C Schaefer, Martina Blank, Angélica R Tamborski, Laura D Guerim, Carla D Bonan, Mònica RM Vianna, and Diogo R Lara. One for all and all for one: the importance of shoaling on behavioral and stress responses in zebrafish. Zebrafish, 10(3):338–342, 2013.

[32] Mirta Galesic, Daniel Barkoczi, Andrew M Berdahl, Dora Biro, Giuseppe Carbone, Ilaria Giannoccaro, Robert L Goldstone, Cleotilde Gonzalez, Anne Kandler, Albert B Kao, et al. Beyond collective intelligence: Collective adaptation. Journal of the Royal Society Interface, 20 (200):20220736, 2023.

[33] L. Lei, R. Escobedo, C. Sire, and G. Theraulaz. Computational and robotic modeling reveal parsimonious combinations of interactions between individuals in schooling fish. PLoS Computational Biology, 16:e1007194, 2020.

[34] Weijia Wang, Ramón Escobedo, Stéphane Sanchez, Clément Sire, Zhangang Han, and Guy Theraulaz. The impact of individual perceptual and cognitive factors on collective states in a data-driven fish school model. PLoS Computational Biology, 18(3):e1009437, 2022.

[35] Roberto E Reis. Check list of the freshwater fishes of South and Central America. Edipucrs, 2003.

[36] Jacques Gautrais, Francesco Ginelli, Richard Fournier, Stéphane Blanco, Marc Soria, Hugues Chaté, and Guy Theraulaz. Deciphering interactions in moving animal groups. PLoS Computational Biology, 8(9):1–11, 09 2012.

[37] Matthew MG Sosna, Colin R Twomey, Joseph Bak-Coleman, Winnie Poel, Bryan C Daniels, Pawel Romanczuk, and Iain D Couzin. Individual and collective encoding of risk in animal groups. Proceedings of the National Academy of Sciences, 116(41):20556–20561, 2019.

[38] Andreu Puy, Elisabet Gimeno, Jordi Torrents, Palina Bartashevich, M Carmen Miguel, Romualdo Pastor-Satorras, and Pawel Romanczuk. Selective social interactions and speed-induced leadership in schooling fish. Proceedings of the National Academy of Sciences, 121(18):e2309733121, 2024.

[39] Kolbjørn Tunstrøm, Yael Katz, Christos C Ioannou, Cristián Huepe, Matthew J Lutz, and Iain D Couzin. Collective states, multistability and transitional behavior in schooling fish. PLoS Computational Biology, 9(2):e1002915, 2013.

[40] Haroldo V Ribeiro, Matthew R Acre, Jacob D Faulkner, Leonardo R da Cunha, Katelyn M Lawson, James J Wamboldt, Marybeth K Brey, Christa M Woodley, and Robin D Calfee. Effects of shady environments on fish collective behavior. Scientific Reports, 12(1):17873, 2022.

[41] Baptiste Lafoux, Paul Bernard, Benjamin Thiria, and Ramiro Godoy-Diana. Confinement-driven state transition and bistability in schooling fish. Physical Review E, 110(3):034613, 2024.

[42] Li Jiang, Luca Giuggioli, Andrea Perna, Ramon Escobedo, Valentin Lecheval, Clément Sire, Zhangang Han, and Guy Theraulaz. Identifying influential neighbors in animal flocking. PLoS Computational Biology, 13(11):e1005822, 2017.

[43] Valentin Lecheval, Li Jiang, Pierre Tichit, Clément Sire, Charlotte K Hemelrijk, and Guy Theraulaz. Social conformity and propagation of information in collective uturns of fish schools. Proceedings of the Royal Society B: Biological Sciences, 285(1877):20180251, 2018.

[44] Benjamin Gallois and Raphaël Candelier. Fasttrack: an open-source software for tracking varying numbers of deformable objects. PLoS Computational Biology, 17(2):e1008697, 2021.

