## Supplementary Information for Experimental evidence of stress-induced critical state in schooling fish for "Experimental evidence of stress-induced critical state in schooling fish"

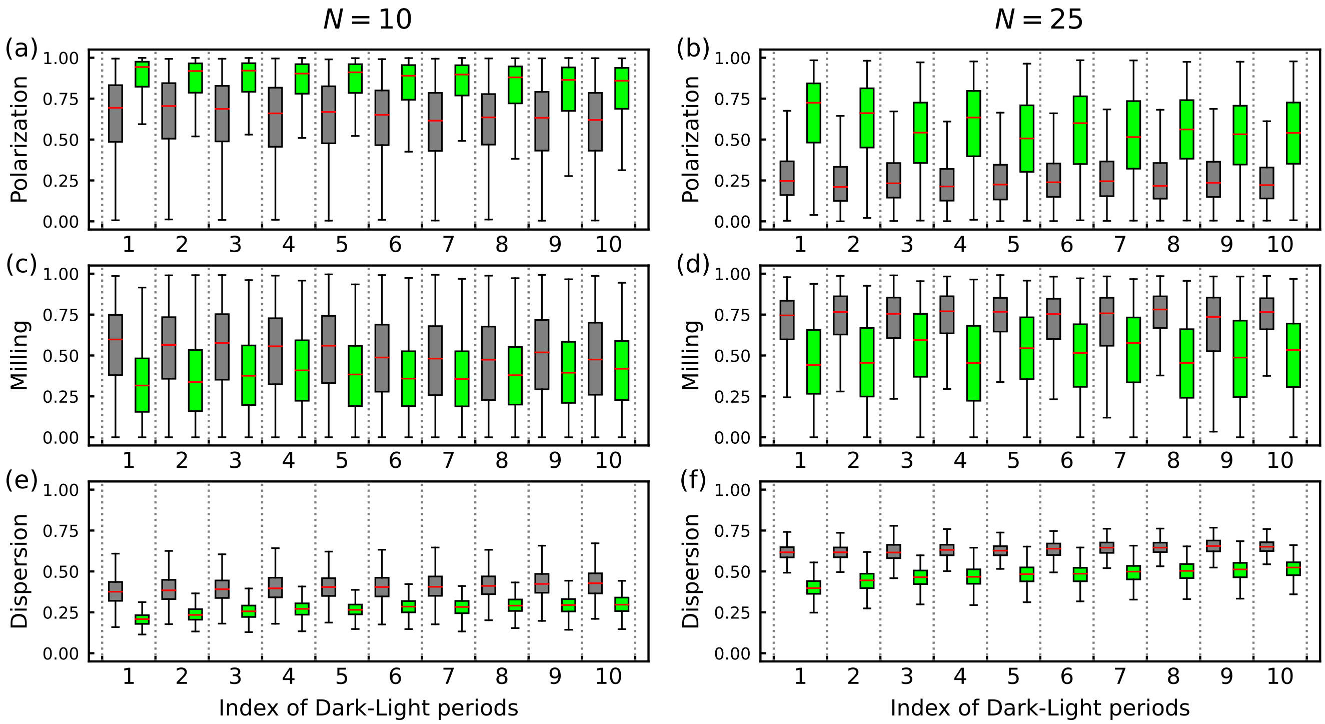


FIG. S1. **Box plots of (a, b) polarization, (c, d) milling, and (e, f) dispersion for $N = 10$ and $N = 25$ in experiments of Stressing condition.** The index of the horizontal axis represents the 10 Dark-Light periods of each experiment in order from left to right, with gray representing the Dark period and green representing the Light period. Each box collects data from all 6 sets of experiments within the corresponding period, the upper and lower whiskers of the box represent the maximum and minimum values, the upper and lower borders of the box represent the third quartile and the first quartile of the data, respectively, and the red line in the box represents the median of the data.

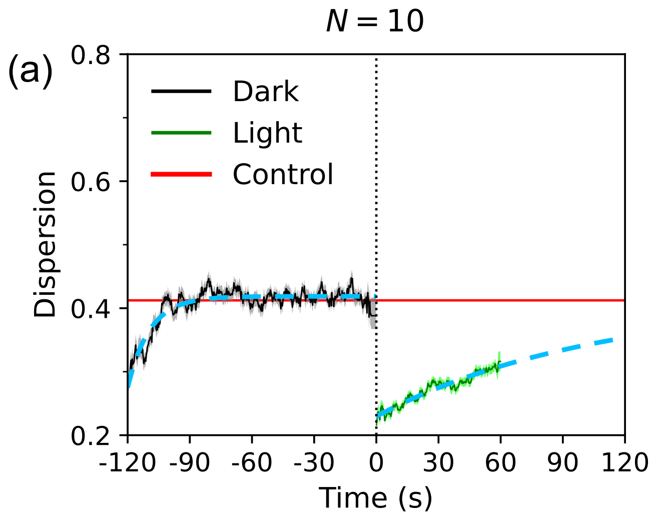


FIG. S2. **Exponential fitting (blue dashed lines) of the time-series of dispersion $\langle D \rangle(t)$ for (a) $N = 10$ group and (b) $N = 25$ group.** The fitting function is of the form $D(t) = C + Ae^{-t/\tau}$, where $C$ is the value of saturation and $\tau$ the relaxation time. In the Light period, $C$ is given by the mean dispersion obtained in the Control condition (red line). In the Dark period, $C$ is obtained by least-squares fitting. The saturation values in the Dark and Light periods are $C = 0.42$ and 0.41, respectively, for $N = 10$, and $C = 0.64$ and 0.69, respectively, for $N = 25$. The relaxation times in the Dark and Light periods are $\tau = 10.70\,\mathrm{s}$ and $105.82\,\mathrm{s}$, respectively, for $N = 10$, and $\tau = 12.73\,\mathrm{s}$ and $95.32\,\mathrm{s}$, respectively, for $N = 25$. The values of $A$ in the Dark and Light periods are $A = -0.14$ and $-0.18$, respectively, for $N = 10$, and $A = -0.17$ and $-0.29$, respectively, for $N = 25$.

| Condition | Group size | Date (y-m-d) | Duration (min) | Usable frames (%) |
|---|---|---|---|---|
| Stressing | N=10 | 2022-08-26 | 29.4 | 84.6 |
| | | 2022-08-26 | 29.3 | 87.8 |
| | | 2022-08-27 | 29.7 | 88.0 |
| | | 2022-08-28 | 29.6 | 91.0 |
| | | 2022-08-28 | 30.0 | 90.0 |
| | | 2022-08-29 | 30.0 | 82.9 |
| | N=25 | 2022-08-26 | 29.4 | 57.0 |
| | | 2022-08-26 | 30.3 | 73.2 |
| | | 2022-08-27 | 29.6 | 71.7 |
| | | 2022-08-28 | 29.4 | 55.2 |
| | | 2022-08-29 | 30.2 | 51.4 |
| | | 2022-08-29 | 29.2 | 68.2 |
| Control | N=10 | 2023-01-03 | 60.0 | 97.3 |
| | | 2023-01-03 | 60.0 | 98.1 |
| | | 2023-01-04 | 60.0 | 98.9 |
| | | 2023-01-04 | 60.0 | 99.0 |
| | N=25 | 2023-01-03 | 60.0 | 94.2 |
| | | 2023-01-03 | 60.0 | 95.8 |
| | | 2023-01-04 | 60.0 | 90.5 |
| | | 2023-01-04 | 60.0 | 92.3 |

TABLE S1. **Summary of experiments.** Includes dates, duration of recorded videos, and proportion of usable video frames after pre-processing for 12 sets of Stressing condition experiments and 8 sets of Control condition experiments.

| Observable \ Condition | | $N = 10$ | | | $N = 25$ | | |
|---|---|---|---|---|---|---|---|
| | | Dark | Light | Control | Dark | Light | Control |
| $D$ | experiment | 0.419±0.081 | 0.281±0.058 | 0.412±0.057 | 0.642±0.043 | 0.492±0.067 | 0.691±0.049 |
| | simulation | 0.419±0.002 | 0.300±0.002 | 0.411±0.003 | 0.642±0.001 | 0.502±0.003 | 0.700±0.003 |
| $P$ | experiment | 0.649±0.218 | 0.827±0.183 | 0.648±0.248 | 0.253±0.157 | 0.552±0.239 | 0.159±0.135 |
| | simulation | 0.633±0.003 | 0.820±0.005 | 0.663±0.004 | 0.260±0.002 | 0.540±0.007 | 0.170±0.002 |
| $M$ | experiment | 0.529±0.255 | 0.396±0.226 | 0.482±0.249 | 0.732±0.194 | 0.490±0.252 | 0.895±0.139 |
| | simulation | 0.525±0.003 | 0.411±0.005 | 0.473±0.005 | 0.716±0.001 | 0.480±0.005 | 0.896±0.001 |

TABLE S2. **Mean values and standard deviations of dispersion, polarization, and milling in experiments and simulations in different light conditions and different group sizes**. The values in the table correspond to the graphical markers shown on each PDF curve in Fig. 3. The last 30 seconds intervals in the Dark and Light period were treated as the steady-state for measurement, and the entire 60 minutes interval in the Control condition was treated as the steady-state.

SM Movie S1: **Group of 10 fish swimming in Stressing condition**. Video excerpt of an experiment with a group of 10 fish swimming in a circular tank of radius 25 cm with an alternation of a period of 2 minutes at 0.5 lx and a period of 1 minute at 25 lx.

SM Movie S2: **Group of 25 fish swimming in Stressing condition**. Video excerpt of an experiment with a group of 25 fish swimming in a circular tank of radius 25 cm with an alternation of a period of 2 minutes at 0.5 lx and a period of 1 minute at 25 lx.

SM Movie S3: **Group of 10 fish swimming in Control condition**. Video excerpt of an experiment with a group of 10 fish swimming in a circular tank of radius 25 cm under constant light intensity at 0.5 lx.

SM Movie S4: **Group of 25 fish swimming in Control condition**. Video excerpt of an experiment with a group of 25 fish swimming in a circular tank of radius 25 cm under constant light intensity at 0.5 lx.

SM Movie S5: **Numerical simulation of the model with a group of 10 fish in Stressing condition (alternating a 2-minutes period at 0.5 lx with a 1-minute period at 25 lx)**. The dots represent individual fish and the attached lines represent the trajectories over a 0.2-second period. The parameter $\gamma_{\mathrm{R}}$ was set to 0.2 for the first two minutes and 0.04 for the last one minute. The variations of $\gamma_{\mathrm{Att}}$ and $\gamma_{\mathrm{Ali}}$ in the simulation refer to Fig. 4 in the main text.

SM Movie S6: **Numerical simulation of the model with a group of 25 fish in Stressing condition (alternating a 2-minutes period at 0.5 lx with a 1-minute period at 25 lx)**. The dots represent individual fish, and the attached lines represent the trajectories over a 0.2-second period. The parameter $\gamma_{\mathrm{R}}$ was set to 0.16 for the first two minutes and 0.34 for the last one minute. The variations of $\gamma_{\mathrm{Att}}$ and $\gamma_{\mathrm{Ali}}$ in the simulation refer to Fig. 4 in the main text.

SM Movie S7: **Numerical simulation of the model with a group of 10 fish in the Control condition (constant light intensity at 0.5 lx)**. The dots represent individual fish, and the attached lines represent the trajectories over a 0.2-second period. The simulation parameters are set to $\gamma_{\mathrm{R}} = 0.24$, $\gamma_{\mathrm{Att}} = 0.023$ and $\gamma_{\mathrm{Ali}} = 0.022$.

SM Movie S8: **Numerical simulation of the model with a group of 25 fish in the Control condition (constant light intensity at 0.5 lx)**. The dots in the video represent individuals, and their attached lines represent trajectories over a 0.2-second period. The simulation parameters are set to $\gamma_{\mathrm{R}} = 0.04$, $\gamma_{\mathrm{Att}} = 0.013$ and $\gamma_{\mathrm{Ali}} = 0.004$.
